# The Alzheimer’s therapeutic Lecanemab induces an amyloid-clearing program in microglia

**DOI:** 10.1101/2025.07.24.666347

**Authors:** Albertini Giulia, Zielonka Magdalena, Cuypers Marie-Lynn, Snellinx An, Xu Ciana, Poovathingal Suresh, Wojno Marta, Davie Kristofer, van Lieshout Veerle, Craessaerts Katleen, Wolfs Leen, Pasciuto Emanuela, Jaspers Tom, Horré Katrien, Serneels Lutgarde, Fiers Mark, Dewilde Maarten, De Strooper Bart

## Abstract

Controversies over anti-amyloid immunotherapies underscore the need to elucidate their mechanisms of action. Here we demonstrate that Lecanemab, a leading anti-Aβ antibody, mediates amyloid clearance by triggering effector functions in the microglia. Using a human microglia xenograft model, we show that Lecanemab significantly reduces Aβ pathology and associated neuritic damage, while neither Fc-inactivated Lecanemab nor microglia deficiency elicit this effect despite intact plaque binding. Single-cell RNA sequencing and spatial transcriptomic analyses reveal that Lecanemab induces a focused transcriptional program that enhances phagocytosis, lysosomal degradation, metabolic reprogramming, interferon gamma genes, and antigen presentation. Finally, we identify SPP1/osteopontin as a major factor induced by Lecanemab treatment and demonstrate its role in promoting Aβ clearance. These findings highlight that effective amyloid removal depends on the engagement of microglia through Fc fragment, providing critical insights for optimizing anti-amyloid therapies in AD.

## Introduction

Lecanemab, an antibody engineered to target soluble amyloid (A)β protofibrils^1^, effectively removes amyloid plaques from the brains of Alzheimer’s disease (AD) patients, slowing cognitive decline by 27%^2^. Although originally developed against peptides containing the rare Artic mutation causally linked to inherited AD^1^, Lecanemab also shows efficacy in sporadic AD cases. Nonetheless, the precise mechanism by which its binding to Aβ oligomers leads to Aβ- plaque clearance compared to other Aβ binding antibodies^3^, remains unclear.

One prevailing hypothesis suggests that plaque clearance is mediated by Fcγ receptor (FcγR) activation of microglia, triggering phagocytosis of Aβ^4–6^. However, direct experimental evidence linking microglia activity to the therapeutic efficacy of Lecanemab is lacking. For instance, while some studies report microglia accumulation around amyloid plaques following immunotherapy^5^, such clustering is also observed without antibody treatment and does not necessarily result in Aβ-plaque removal. Moreover, FcγR activation can induce a pro-inflammatory response with the release of cytokines and other toxic mediators^7,8^, potentially mitigating the benefits of Aβ clearance. Alternative FcγR-independent mechanisms of plaque removal have also been widely proposed^9–12^. Consequently, the exact mechanism by which Lecanemab clears amyloid plaques remains unresolved.

To uncover the mechanisms driving antibody-induced amyloid clearance, we generated Lecanemab^13^ alongside a human IgG1 variant designed to abolish Fc-mediated effector functions^14,15^, Lecanemab LALA-PG. We leveraged our advanced human microglia xenotransplantation model for AD^16,17^. Using Rag2^tm1.1Flv^; Csf1^tm1(CSF1)Flv^; Il2rg^tm1.1Flv^; App^tm3.1Tcs^; Csf1R^em1Bdes^ mice, named hereafter as App^NL-G-F^ Csf1r^ΔFIRE/ΔFIRE^ mice, which lack endogenous microglia^17,18^, enabled us to assess these clinically relevant human antibodies directly. Importantly we corroborated our observations in fully immune-competent mice treated with murine analogues of Lecanemab. Our work probes a critical conundrum: why do microglia— although strongly activated in the presence of amyloid plaques—fail to clear these deposits? We hypothesize that antibody engagement unveils latent, transformative mechanisms, endowing microglia with an unexpected capacity for plaque clearance. Our work suggests that antibody treatment reprograms microglial function, providing insights into protective roles of these cells while also opening new avenues to understand and ultimately harness these changes for therapeutic benefit.

## Results

### Lecanemab binds to amyloid plaques

By 4 months of age, xenografted human microglia had efficiently colonized the mouse brain (**Extended Data Fig. 1a-b**) and, by 6 months, they are fully able to mount amyloid responses which strongly resemble the ones observed in AD patients (as previously characterized in^16^) (**Extended Data Fig. 1c**). Starting from 4 months of age, we administered weekly intraperitoneal injections of 10 mg/kg Lecanemab or Lecanemab LALA-PG^19^. After eight weeks of treatment, we analyzed the distribution of the human antibodies in the brain parenchyma (**Fig. 1a-b**). Strikingly, while only sparse antibody signals were detected in Lecanemab-treated mice (notably within human CD45⁺ microglia surfaces, as reconstructed in **Supplementary Movie 1**), Lecanemab LALA-PG strongly accumulated on Aβ plaques. This accumulation is already evident after 2 weeks of treatment (**Fig. 1c**). This data suggests that mutating the Fc fragment to abolish Fc-based effector functions prevents uptake of Lecanemab into microglia. Additionally, our findings demonstrate that Lecanemab binds to plaques, challenging the common assumption that it is specific to oligomers. These remarkable findings led us to investigate the potential changes these antibodies might induce in the human microglia surrounding the Aβ plaques^20,21^.

**Figure 1.**
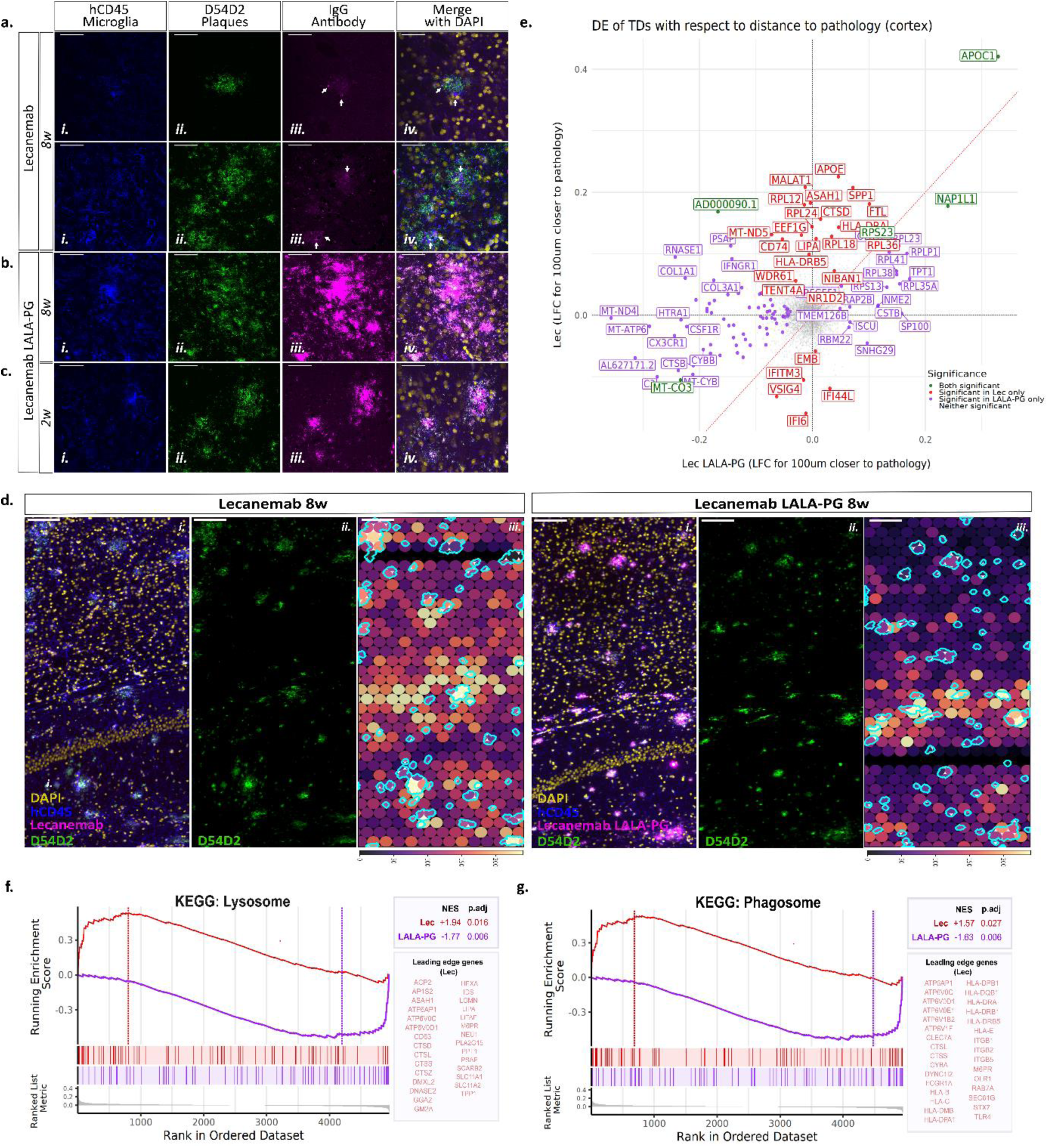
Lecanemab drives strong transcriptional changes in human microglia associated to Aβ plaques. **a-c.** Representative high-magnification confocal z-stacks of (*i*) CD45 (human microglia, blue), (*ii*) D54D2 (Aβ, green), (*iii*) IgG (human antibody, magenta) and (*iv*) a merged view with DAPI-stained nuclei (yellow). These high resolution z-stacks are used to show the colocalization between D54D2 and IgG, as well as the internalization of Lecanemab within the microglia. Scale bar: 50 µm. **a.** After 8 weeks of Lecanemab administration, human IgG is detected in the brain parenchyma, where it associates with Aβ plaques and is internalized by human microglia (arrows). **b.** Notably, after 8 weeks of Lecanemab LALA-PG administration, IgG exhibits strong accumulation on D54D2+ Aβ plaques, indicating that a functional Fc fragment is necessary for uptake in microglia. **c.** This accumulation is already apparent as early as 2 weeks after treatment in Lecanemab LALA-PG–treated mice. **d.** Representative large-field images of the Nova-ST data coupled with immunofluorescence workflow for Lecanemab- (upper panel) and Lecanemab LALA-PG- (lower panels) treated mice. Immunofluorescence was performed to visualize human microglia (CD45^+^, blue), Aβ plaques (D54D2^+^, green), Lecanemab or Lecanemab LALA-PG (magenta) and DAPI-stained nuclei (yellow). D54D2 signal was used to define the plaque regions (outlined in cyan). Panels (*i*) show a merged view, panels (ii) D54D2, panels (iii) show spatial transcriptomic TDs (spots binned into hexbins with a diameter of 40 µm) overlayed with plaque ROIs. TDs are colored based on their relative expression of human transcripts (purple: low expression, yellow: high expression). Scale bar: 100 µm. **e.** Quadrant plot showing the log2 fold changes (LFCs) of genes in Lecanemab- (x axis) and Lecanemab LALA-PG- (y axis) treated TDs with respect to their distance to plaques in the Nova-ST dataset. A positive LFC indicates up-regulation in proximity to plaques. Red: genes significant in Lecanemab microglia only; purple: genes significant in Lecanemab LALA-PG microglia only; green: genes significant in both comparisons; and gray: genes not significant in either comparison. LFCs were calculated using edgeR’s quasi-likelihood F-test; p values were adjusted with BH correction (p.adj < 0.05). **f-g**. From the differential gene expression analysis in (**e**), we performed gene set enrichment analysis (GSEA) to further explore shifts in the microglia phenotype following antibody treatment. We saw a significant positive enrichment of phagosome (**f**) and lysosome (**g**) genes in Lecanemab TDs near Aβ plaques (red), whereas such enrichment was not observed in Lecanemab LALA-PG TDs (purple). The vertical dotted lines indicate the enrichment score (ES). Vertical tick marks along the x axis show the location of individual genes in the gene set within the FC-ranked gene list. The Normalized Enrichment Score (NES), p.adj value, and the leading-edge genes are shown for the GSEA performed on the Lecanemab and Lecanemab LALA-PG TDs.

### Spatial transcriptomics reveals Lecanemab-driven enhancement of microglial phagocytic and lysosomal pathways

We employed Nova-ST^22^, a recently developed technique based on Illumina NovaSeq flow cells, which enabled us to combine unbiased high-resolution spatial transcriptomics with immunofluorescence of amyloid plaques (positive for the anti-Aβ antibody D54D2) on the same tissue section (**Fig. 1d**, **Extended data Fig. 2**). Since only the xenotransplanted cells are of human origin, all human-derived reads can be exclusively attributed to microglia^23^.

We first converted the raw spatial expression matrix, binning spots into hexbin pseudo-spots with a width, and center-to-center distance of 40 μm in diameter (tissue domains, TDs) and retained only those with at least 30 human transcripts for further analysis. A total of 32,568 TDs were obtained across the cortical regions of the 4 samples, with 17,186 bins in the Lecanemab treated samples and 15,382 from Lecanemab LALA-PG (**Extended Data Fig. 2a-b**). Each TD captured an average of 61.3 human genes and 74.4 UMI (**Extended Data Fig. 2c**). We next performed a continuous differential expression analysis of cortical bins based on their distance from D54D2^+^ Aβ plaques, revealing that the expression of several genes significantly increases in function of proximity to plaques in the Lecanemab treated mice, including *APOE, CTSD, SPP1, CD74* and other genes associated to antigen presentation (**Fig. 1e**). Notably, Gene Set Enrichment Analysis (GSEA) indicated a significant increase in the expression of pathways related to the phagosome (**Fig. 1f**, **Extended Data Fig. 2d**) and lysosome (**Fig. 1g**, **Extended Data Fig. 2d**), specifically in the Lecanemab-treated TDs. Interestingly, these gene sets were significantly down-regulated in Lecanemab LALA-PG TDs in proximity to pathology (**Fig. 1f-g**, **Extended Data Fig. 2e**), suggesting that the Fc fragment is critical for the activation of these pathways in microglia near plaques. That said, we detected distinct transcriptomic effects induced by the Lecanemab LALA-PG treatment (**Extended Data Fig. 2e-f**), suggesting that the accumulation of non-functional antibody also modulates microglia activity near plaques. This data prompted further investigation into how the observed enhancements in phagocytosis and lysosomal functions by Lecanemab might impact plaque load.

### Lecanemab attenuates Aβ pathology via Fc-Mediated Microglial phagocytosis

To assess the impact of Fc-mediated phagocytosis on Aβ plaque load, we treated a cohort of xenotransplanted App^NL-G-F^ Csf1r^ΔFIRE/ΔFIRE^ mice with Lecanemab, Lecanemab LALA-PG, or human IgG1 control for 8 weeks (**Fig. 2a**). After completion of the treatment (24 hours following the last injection) we harvested the brains (**Fig. 2b**), and we used immunohistochemistry to analyze the impact of treatment on amyloid pathology. We quantified both β-sheeted amyloid aggregates using X34 (**Fig. 2c**), and Aβ-peptides using a specific antibody (82E1) (**Fig. 2d**). Lecanemab treatment significantly reduced plaque area compared to IgG1 or Lecanemab LALA-PG. Interestingly, when analyzing X34^+^ plaque distribution, we observed that Lecanemab had the most pronounced effect on smaller plaques (**Fig. 2e**). Histological data were corroborated by MSD ELISA, which revealed significantly reduced guanidine-extractable (insoluble) Aβ42 levels in Lecanemab-treated mice compared to those treated with IgG1 or Lecanemab LALA-PG (**Fig. 2f**). Notably, Aβ42 is the predominant Aβ species in App^NL-G-F^ mice^24^. A significant reduction in insoluble Aβ38 levels was also observed in Lecanemab-treated mice compared to IgG1-treated mice (**Fig. 2f**), while soluble Aβ38 levels were significantly reduced in Lecanemab-treated mice compared to those treated with Lecanemab LALA-PG (**Fig. 2g**). No changes were observed in insoluble Aβ40 levels (**Fig. 2f-g**). Our App^NL-G-F^ Csf1r^ΔFIRE/ΔFIRE^ mice lack a mature adaptive immune system, to allow colonization of the human brain with xenografted human microglia. To demonstrate that Lecanemab effect on plaque load is not hampered by the presence of adaptive immune cells, we repeated the experiments in a cohort of immunocompetent App^NL-G-F^ mice treated with mAb158 (a murine version of Lecanemab), its engineered LALA-PG variant, and a control mouse IgG2a (**Fig. 2h**). After eight weeks of treatment, mAb158 significantly decreased Aβ levels, as measured by MSD ELISA (**Fig. 2i-j**), whereas the LALA-PG mutation abolished antibody mediated Aβ clearance in this model with an intact murine immune system. Finally, we demonstrated the essential role of microglia as App^NL-G-F^ Csf1r^ΔFIRE/ΔFIRE^ mice that did not receive the xenotransplantation (**Fig. 2k**) showed no impact from Lecanemab on plaque load (**Fig. 2l-p**). Collectively, these findings demonstrate that Lecanemab attenuates Aβ pathology *in vivo* through Fc-mediated microglial effector functions.

**Figure 2.**
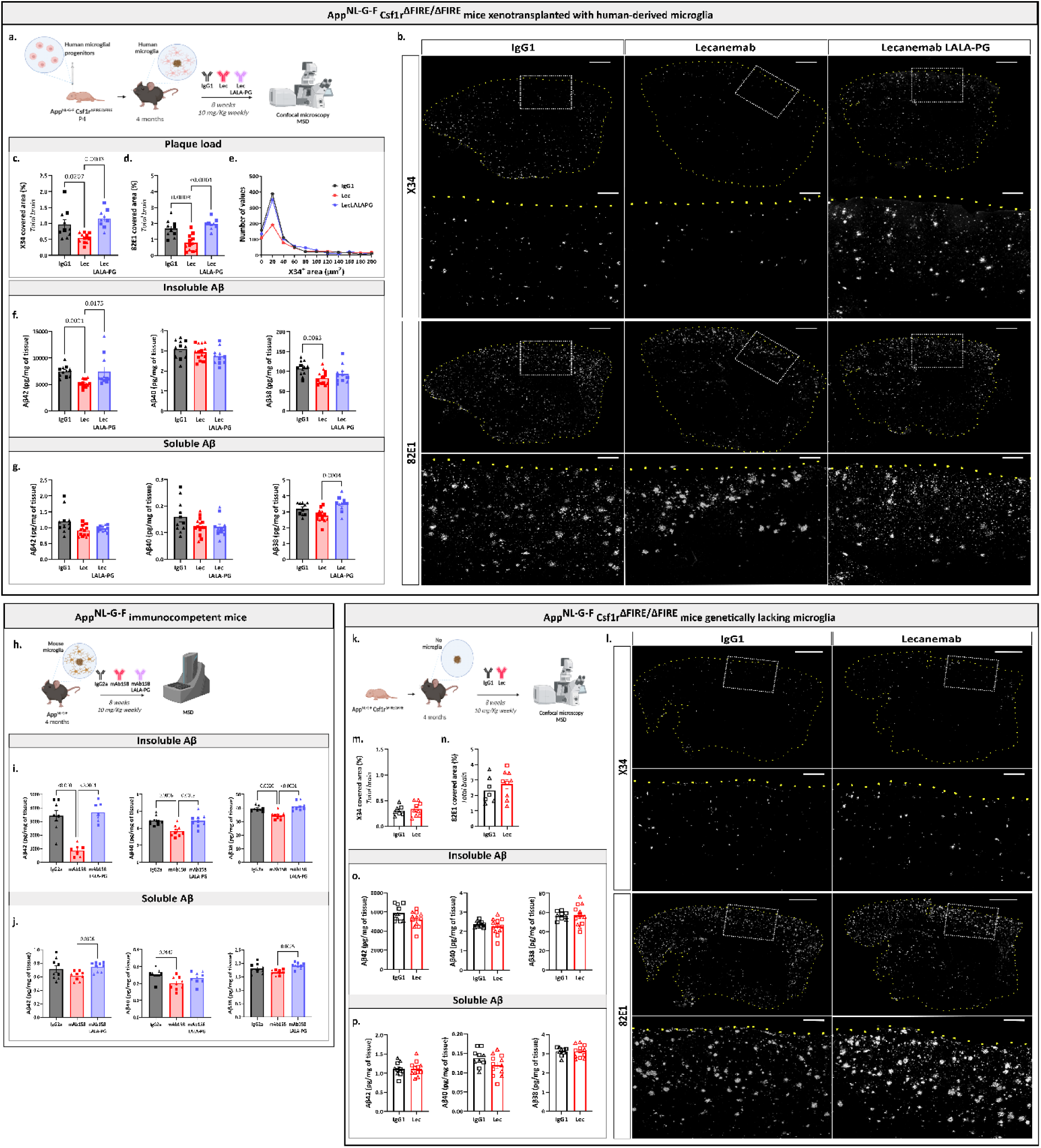
Lecanemab alleviates Aβ pathology by triggering effector functions in the microglia. **a.** App^NL-G-F^ Csf1r^ΔFIRE/ΔFIRE^ mice were xenotransplanted at P4 with human-derived microglial progenitors differentiated *in vitro*. Starting from 4 months of age, mice were treated for 8 weeks with IgG1, Lecanemab or Lecanemab LALA-PG (10 mg/kg, weekly i.p. injections) and sacrificed for subsequent analysis. **b.** Representative confocal images of plaques stained with X34 or 82E1 in sagittal brain sections from mice treated with the indicated antibodies; scale bar: 1 mm; inset: 250 µm. **c-d.** Quantification of X34 (**c**) and 82E1 (**d**) area expressed as percentage of the total section area; (**c**) Kruskal-Wallis test (p = 0.0003) and (**d**) one-way ANOVA (p < 0.0001). **e.** Distribution of X34^+^ plaques based on their area (x-axis) shows- that Lecanemab mainly affects small plaques; Anderson-Darling test, p < 0.0001 (IgG1 vs Lecanemab, p < 0.0003; Lecanemab vs Lecanemab LALA-PG, p < 0.0003; IgG1 vs Lecanemab LALA-PG: n.s.; Kolmogorov-Smirnov test). **f.** MSD ELISA of Aβ42 (p = 0.0001, Kruskal-Wallis test followed by Dunn’s multiple comparison test), Aβ40 (n.s., one-way ANOVA) and Aβ38 (p = 0.0117, one-way ANOVA followed by Bonferroni’s multiple comparisons test) levels in insoluble (GuHCl extractable) brain extracts. **g.** MSD ELISA of Aβ42 (p = 0.0505, Kruskal-Wallis test), Aβ40 (n.s., one-way ANOVA) and Aβ38 (p = 0.0005, one-way ANOVA followed by Bonferroni’s multiple comparisons test) levels in soluble (T-PER buffer extractable) brain extracts. **h.** A cohort of immunocompetent App^NL-G-F^ mice was used to assess the impact of mAb158 (a murine version of Lecanemab), its engineered LALA-PG variant, and a control mouse IgG2a on Aβ levels by MSD. **i.** MSD ELISA of Aβ42 (p < 0.0001, one-way ANOVA followed by Bonferroni’s multiple comparisons test), Aβ40 (p = 0.0005, one-way ANOVA followed by Bonferroni’s multiple comparisons test) and Aβ38 (p < 0.0001, one-way ANOVA followed by Bonferroni’s multiple comparisons test) levels in insoluble brain extracts. **j.** MSD ELISA of Aβ42 (p = 0.0355, one-way ANOVA followed by Bonferroni’s multiple comparisons test), Aβ40 (p = 0.0026, one-way ANOVA followed by Bonferroni’s multiple comparisons test) and Aβ38 (p = 0.0032, one-way ANOVA followed by Bonferroni’s multiple comparisons test) levels in soluble brain extracts. **k.** A cohort of App^NL-G-F^ Csf1r^ΔFIRE/ΔFIRE^ mice was not xenotransplanted and used to assess if Lecanemab alters Aβ pathology in the absence of microglia. **l.** Representative confocal images for X34 and 82E1 in sagittal brain sections from App^NL-G-F^ Csf1r^ΔFIRE/ΔFIRE^ mice treated with IgG1 or Lecanemab; scale bars: 1 mm, inset: 250 µm. **m-n.** Quantification of X34 and 82E1 areas (unpaired t tests, n.s.) expressed as percentage of the section area. **o.** MSD ELISA of Aβ42, Aβ40 and Aβ38 levels in insoluble brain extracts (unpaired t tests, n.s.). **p.** MSD ELISA of Aβ42, Aβ40 and Aβ38 levels in soluble brain extracts (unpaired t tests, n.s.). Mean ± SEM shown for each group and points represent individual animals. Square symbols: males; triangle: females.

We next evaluated the impact of Lecanemab on microglia phagocytosis of Aβ fibrils. In an *in vitro* assay using cryosections from App^NL-G-F^ mouse brains (**Extended Data Fig. 3a**), sections were incubated for 1 hour with control IgG1, Lecanemab, or Lecanemab LALA-PG^14^. Subsequently human-derived microglial cells^16^ were added and after three days, amyloid plaque area was quantified using the pan-Aβ antibody 82E1 (**Extended Data Fig. 3b**). Lecanemab treatment sections exhibited a significantly reduced Aβ-plaque load relative to those exposed to control IgG1 or Lecanemab LALA-PG (**Extended Data Fig. 3c**). Notably, in the absence of microglia, no plaque clearance was observed (**Extended Data Fig. 3d**) confirming that Lecanemab facilitates microglia-mediated Aβ-clearance *in vitro*. To validate these findings *in vivo*, a cohort of App^NL-G-F^ mice xenotransplanted with human microglia was treated with either Lecanemab or IgG1 from 6 to 8 months of age. Following treatment, mice received an intraperitoneal injection of methoxy-XO4, a fluorescent probe that crosses the blood-brain barrier to stain Aβ^25^. Three hours post-injection, human CD45⁺microglia were harvested and the proportion of methoxy-XO4⁺ cells was quantified by flow cytometry (**Fig. 3a**, **Extended Data Fig. 3f**). Lecanemab treatment resulted in a significant increase in Aβ uptake by hCD45⁺ microglia, particularly within the CD68^high^ subset (**Fig. 3b-c**).

**Figure 3.**
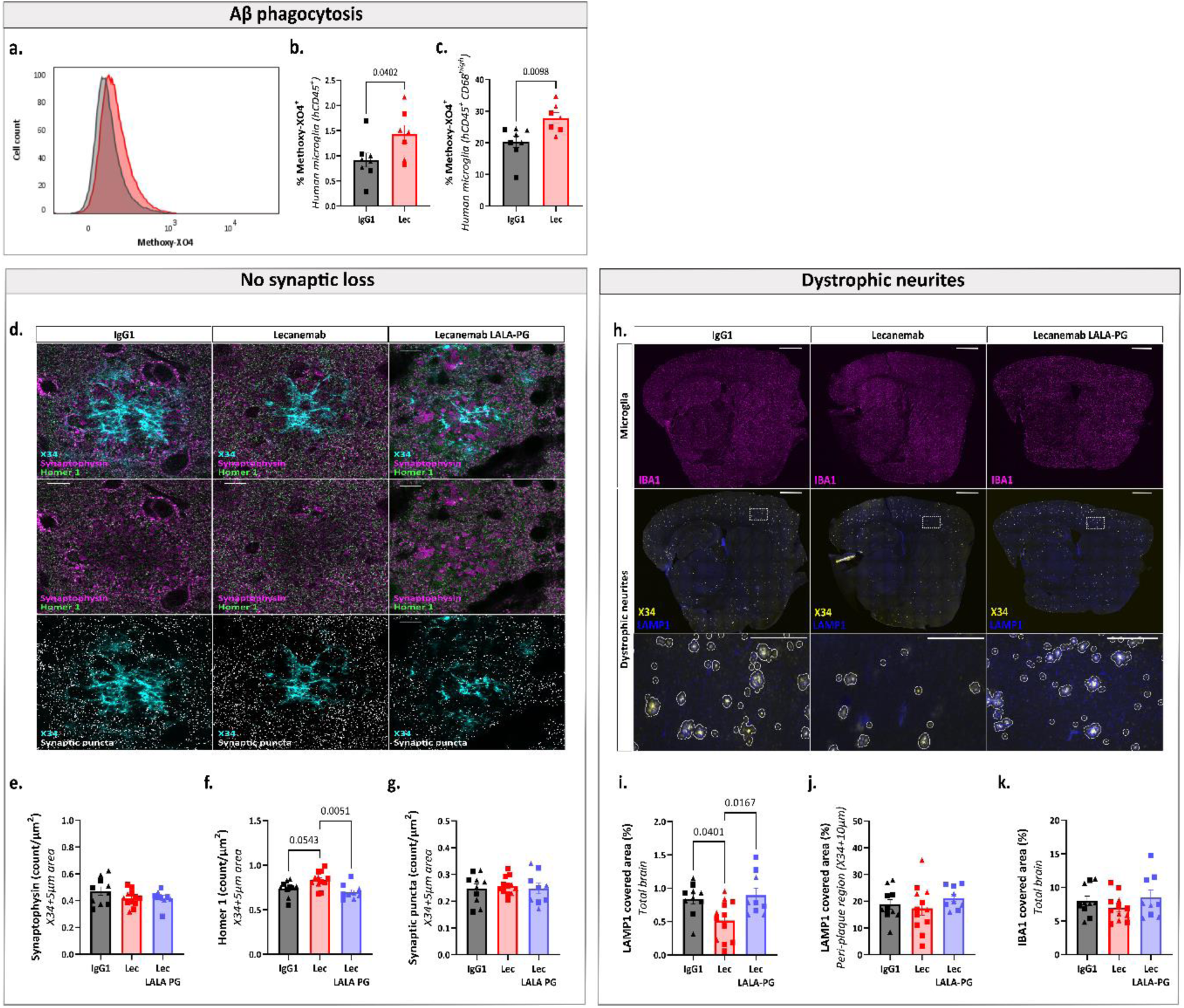
Lecanemab induces phagocytosis of Aβ and alleviates downstream consequences of Aβ pathology through Fc-mediated microglial effector functions. **a.** Flow cytometry analysis of Methoxy-X04 positive cells within the hCD45^+^ population (microglia) after 8 weeks of Lecanemab (red) or IgG1 (black) treatment. The x-axis represents Methoxy-X04 fluorescence intensity, while the y-axis shows the number of cells. The overlaid histograms (black and red) highlight differences in Methoxy-X04 staining levels. **b.** Percentage of XO4+ microglia (CD45^+^) isolated from IgG1 or Lecanemab treated mice, unpaired t tests. **c.** Percentage of XO4^+^ microglia (CD45^+^, CD68^high^) isolated from IgG1 or Lecanemab treated mice, unpaired t tests. **d.** Representative super-resolution confocal images of synaptic loss surrounding X34^+^ plaques in IgG1-, Lecanemab- and Lecanemab LALA-PG-treated xenotransplanted mice. Synaptic puncta were defined as synaptophysin (magenta) and homer 1 (green) immunoreactive puncta (white) around X34^+^ plaques (cyan). Scale bar: 10 µm. **e-g**. Quantification of synaptophysin (one-way ANOVA, n.s.), homer 1 (one-way ANOVA, p = 0.044) and synaptic puncta (one-way ANOVA, n.s.), in peri-plaque area (defined as 5 µm from the X34 edges). **h.** Representative confocal images for IBA1 (magenta), X34 (yellow) and LAMP1 (blue) in sagittal brain sections from App^NL-G-F^ Csf1r^ΔFIRE/ΔFIRE^ mice xenotransplanted with human-derived microglia and treated with IgG1, Lecanemab and Lecanemab LALA-PG; scale bars: 1 mm and 250 µm in the inset. The area of LAMP1 was assessed in the 10 µm-rings surrounding X34 (peri-plaque region) and then divided by the area of the brain section or the peri-plaque area. **i-j.** Quantification of LAMP1 area expressed as percentage of the total brain section (**i**) and the peri-plaque area (**j**); (**i**) one-way ANOVA (p = 0.0091) and (**j**) one-way ANOVA (n.s.). **k.** Quantification of IBA1 area expressed as percentage of the total brain section (Kruskal-Wallis test, n.s.). Mean ± SEM shown for each group and points represent individual animals. Square symbols: males; triangle: females.

Given that increased phagocytosis may be detrimental if non-specific, for instance, leading to removal of synapses in the proximity of plaques^9^, we assessed the density of the presynaptic marker Synaptophysin, the post-synaptic marker Homer 1, and their overlap (synaptic puncta) in the peri-plaque area (defined as 5 µm from the X34 edges, **Fig. 3d** ). Synaptophysin (**Fig. 3e**) or synaptic puncta densities (**Fig. 3g**) remained unchanged, and we found a small but significant increase in post-synaptic density compared to IgG1- and Lecanemab LALA-PG-treated animals (**Fig. 3f**). These findings suggest that Lecanemab-induced phagocytosis is specific to Aβ. We further investigated the downstream effects of reduced plaque load on Aβ- related pathologies. Lecanemab treatment significantly decreased neuritic pathology, as indicated by LAMP1 staining across the total brain area (**Fig. 3h-i**). However, the ratio of LAMP1 positive area to amyloid plaque load was unchanged (**Fig. 3j**), suggesting that the reduction in dystrophic neurites is an indirect consequence of decreased plaque load.

### Lecanemab remodels microglial transcriptome to activate clearance pathways

Microglia clear amyloid plaques effectively only when Lecanemab is present, suggesting that Fc-engagement modifies their functionality. To investigate the mechanisms underlying plaque clearance with greater resolution, we performed single-cell transcriptomic profiling of human microglia using the 10X Genomics platform, achieving deeper gene expression coverage than spatial transcriptomics. Because our data indicated that Lecanemab LALA-PG strongly accumulates on Aβ-plaques affecting microglia responses (**Fig. 1b-c**), we compared microglia treated with Lecanemab to those treated with a control IgG1. Following quality control and removal of macrophages (**Extended Data Fig. 4a-d**), a differential gene expression analysis between IgG1- and Lecanemab-treated human microglia revealed approximately 300 significantly differentially expressed genes (**Fig. 4a**). Consistent with our spatial transcriptomic results, upregulated genes were enriched for terms associated with phagosome pathway and antigen presentation (**Extended Data Fig. 4e**). Additionally, we identified genes enriched for processes related to interferon response, metabolism, and the unfolded protein response (**Extended Data Fig. 4e**).

**Figure 4.**
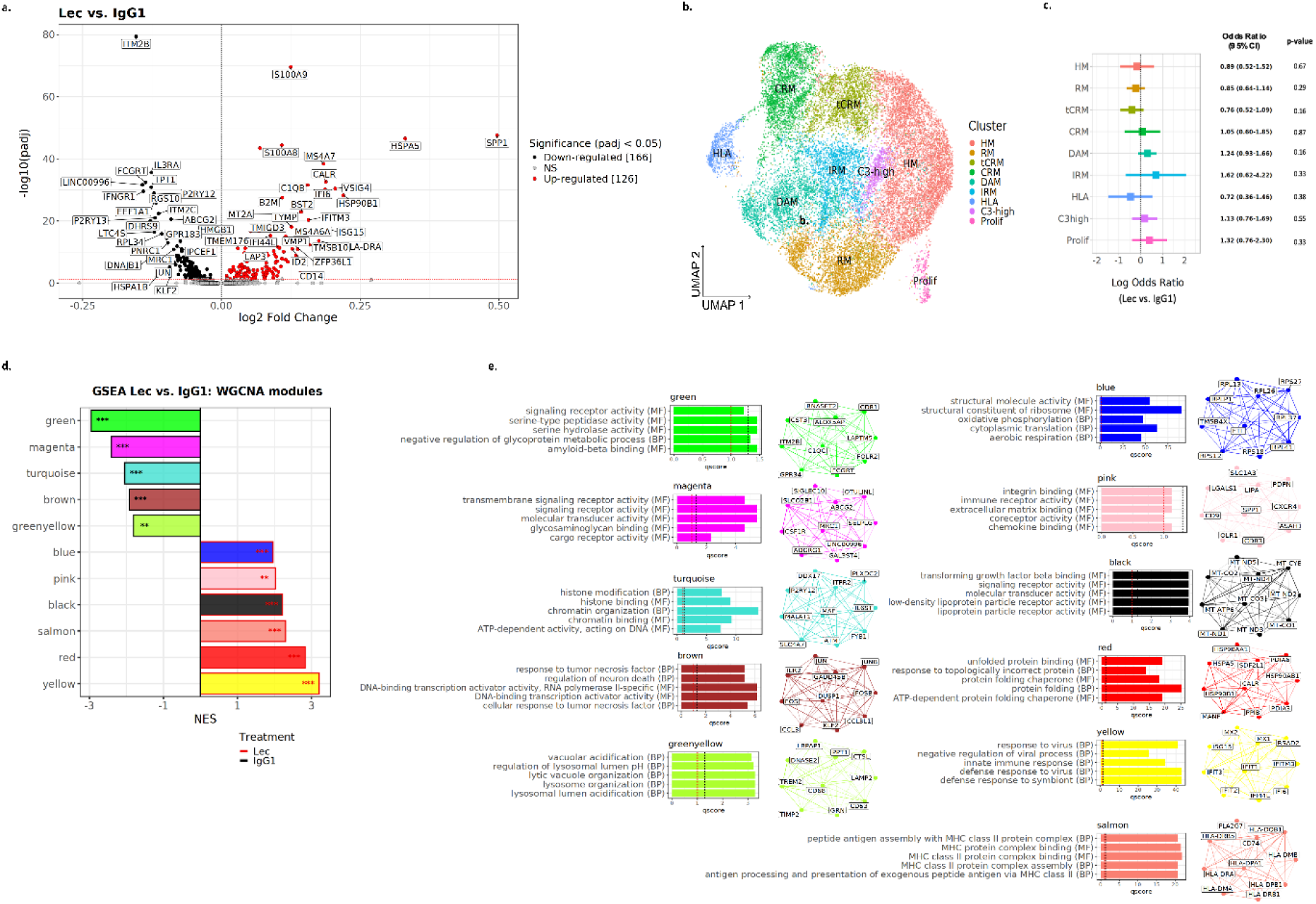
Transcriptomic changes in human microglia treated with Lecanemab. **a**. Volcano plots showing a gene expression comparison between IgG1- and Lecanemab-treated human microglia. The number of significant genes per condition is reported in brackets. Adjusted p-value threshold < 0.05 (Wilcoxon rank-sum test, p-values adjusted with Bonferroni correction based on the total number of genes in the dataset, NS = not significant). **b.** UMAP plot visualizing 22,420 cells (10,850 iGg1, 11,570 Lecanemab) human microglial cells after removal of macrophages. Cells are colored according to clusters identified. The assignment of different clusters to distinct cell types or states is based on previous experimental data from our laboratory^14^. **c.** Odds ratio and 95% confidence interval for the differential abundance of cell states between Lecanemab- and IgG1-treated group using MASC analysis. No significant changes in cell states proportion are detected between IgG1 and Lecanemab-treated microglia. **d.** NES of significantly enriched (p.adj < 0.05) Weighted Gene Co-expression Network Analysis (WGCNA) modules between IgG1 and Lecanemab treated cells, as identified by GSEA (*: p.adj < 0.05, **: p.adj < 0.01, ***: p.adj < 0.001). **e.** Functional annotations based on Gene Ontology (GO) pathway analyses (MF: molecular function and BP: biological process; enrichment for a given ontology is shown by q score: red line p.adj<0.1, black line p.adj<0.05) and top 10 hub genes by module kME.

While we observed changes in expression levels, their magnitude is relatively subtle. Furthermore, Lecanemab’s efficacy does not appear to be driven by the induction of specific cell states, which ultimately reflect broad transcriptional changes, such as DAM/HLA state (**Fig. 4b-c**, **Extended Data Fig. 4f**). This insight prompted us to investigate whether the gene set induced by Lecanemab is more specifically targeted. To do so, we performed weighted gene co-expression network analysis (WGCNA) and identified 14 modules of co-expressed genes (**Extended Data Fig. 5a-c**, **Table 1**). Gene set enrichment analysis (GSEA) revealed that five modules were significantly downregulated and six were significantly upregulated following Lecanemab treatment (**Fig. 4d**). Among the upregulated modules (**Fig. 4e**, **Table 2**) the yellow module was enriched for interferon genes, the red module for unfolded protein and protein folding genes, the salmon module for antigen presentation genes, the black module for the mitochondrial and immune signaling genes, and the blue module for metabolism genes, particularly those involved in oxidative phosphorylation and aerobic respiration. The pink module, although not associated with a specific functional signature in Gene Ontology (GO) databases, was enriched for *SPP1*, *LGALS1, CTSD* and *ITGAX,* core genes of protective axon-tract-associated microglia (ATM) and proliferative-region-associated microglia (PAM) identified during early microglia development^26,27^ (**Fig. 5a**, **Table 3**). Notably, *SPP1* (encoding osteopontin/OPN), is also the most strongly upregulated gene in both our scRNA-seq and spatial differential expression analyses. *SPP1* is implicated in phagocytosis^26,28^ and has been linked to a protective response in developing microglia^26^. Although *SPP1* is a well-established marker of disease-associated microglia (DAM/HLA)^16^, which typically accumulate around amyloid plaques without clearing them, our findings suggest that Lecanemab induces even higher expression of this gene and might activate clearance programs that are not fully engaged in DAM/HLA.

**Figure 5.**
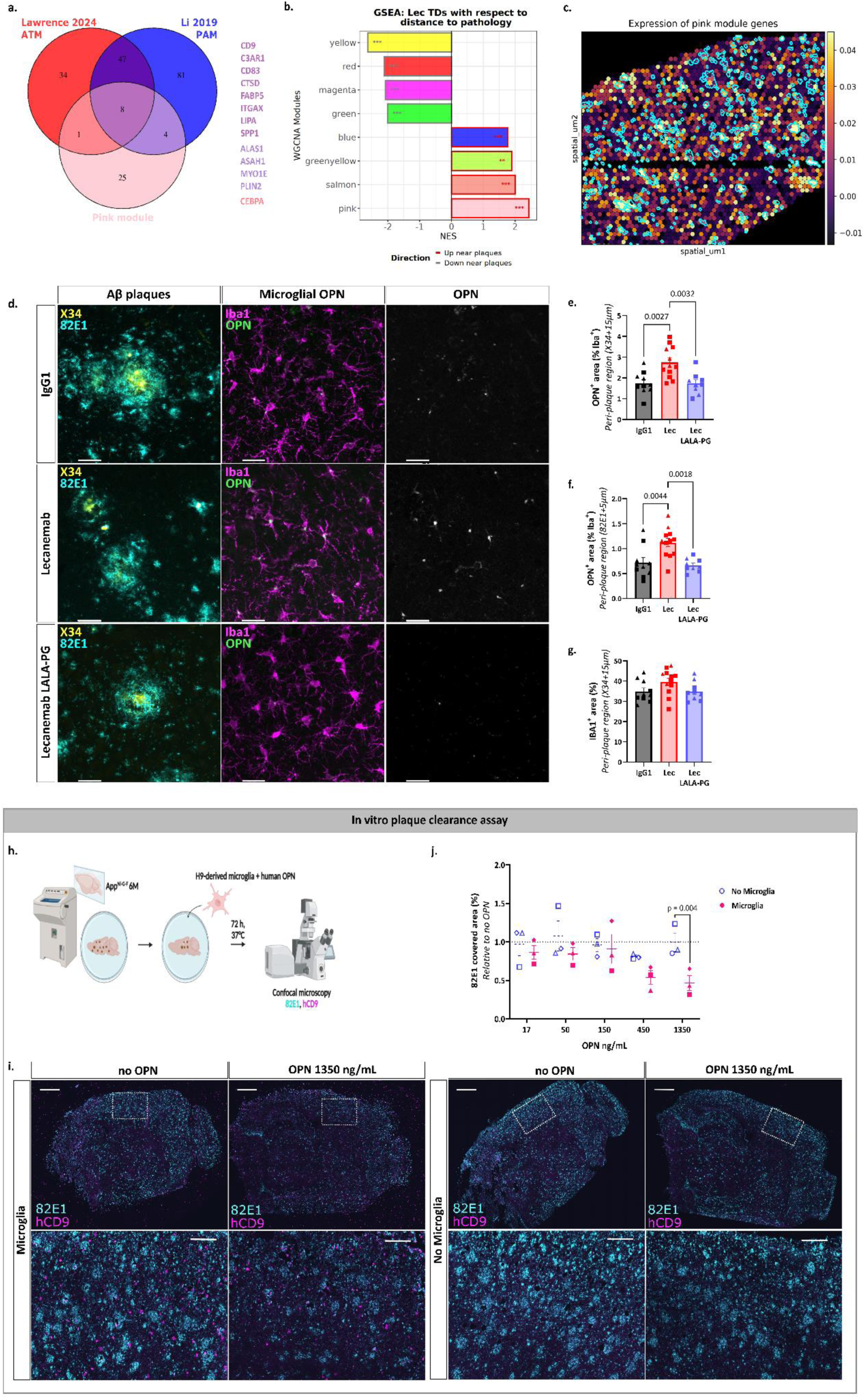
Osteopontin/SPP1, one of the main factors induced by Lecanemab treatment, promotes Aβ clearance. **a.** Venn diagram illustrating the overlap between the ATM (red)^26^, PAM (blue)^27^, and the pink module gene sets. Eight genes are common to all three sets, one gene is shared exclusively between the ATM and the pink module, while four genes are shared exclusively between the PAM and the pink module. The list of shared genes is displayed alongside the diagram. **b.** Normalized enrichment score (NES) of Weighted Gene Co-expression Network Analysis (WGCNA) modules in Lecanemab tissue domains (TDs, 40 μm hexbin pseudo-spots) with respect of distance to pathology, as identified by Gene Set Enrichment Analysis (GSEA) performed on the differential expression analysis in Fig. 1e (**: p.adj < 0.01, ***: p.adj < 0.001). **c.** Cortical TDs in Lecanemab-treated mice colored based on their relative expression of genes belonging to the pink module (purple: low expression, yellow: high expression) and overlayed with plaque ROIs (white). Enrichment scores were obtained using scanpy’s *score_genes()* function. Note the significant enrichment of the module in TDs close to Aβ plaques, as quantified in (**b**). **d.** Representative confocal images of cortical X34+ and 82E1+ plaques surrounded by OPN+ microglia in IgG1-, Lecanemab- and Lecanemab LALA-PG-treated mice. **e-f**. Quantification of the area of OPN+ area within IBA1+ cells around X34+ (**e**) and 82E1+ plaques (**f**) in IgG1- and Lecanemab-treated mice); (**e**) one-way ANOVA (p = 0.0008) and (***j***) one-way ANOVA (p = 0.0007). **g.** Quantification of the area covered by IBA1+ cells around X34+ plaques; one-way ANOVA (n.s.). **h.** Schematic representation of the *in vitro* plaque clearance assay paradigm used to study Aβ clearance in response to OPN stimulation. Human-derived microglial cells were plated onto sagittal cryosections from 6-month-old App^NL-G-F^ mice, followed by treatment with increasing concentrations of human OPN (17, 50, 150, 450, and 1350 ng/mL). After three days, Aβ plaque coverage was quantified using the pan-Aβ antibody 82E1. **i.** Representative confocal images of 82E1 (Aβ, cyan) and CD9 (human microglia, magenta) immunoreactivity in App^NL-G-F^ brain cryosections after OPN stimulation. Scale bar: 1 mm; inset: 200 µm. **j.** Quantification of 82E1 covered area (relative to no OPN) in sections plated with or without human microglia; modified Chi-squared method (p = 0.03). Graphs show mean±SEM and points represent individual animals (**e-g**) or independent experiments being each the average of 1-4 cryosections (**j**). Square symbols: males; triangles: females.

We then used our functional WGCNA modules generated on the scRNA-seq data to interpret changes in the spatial transcriptomic data relative to plaque proximity. Notably, two key GSEA-identified modules, the yellow module (interferon-related) and the red module (unfolded protein response), do not seem to be associated to plaques, although significantly enriched in the Lecanemab TDs compared to the Lecanemab LALA-PG TDs (**Extended Data** Fig. 5e). On the other hand, the pink module (enriched for *SPP1*) was the most significantly enriched in plaque associated TDs (**Fig. 5b-c**), followed by modules associated with antigen presentation (salmon), lysosome (green-yellow) and metabolism (blue). Consistent with our transcriptomic data, OPN expression was elevated in IBA1^+^ microglia surrounding X34^+^ and 82E1^+^ plaques in Lecanemab-treated mice compared to those treated with IgG1 or Lecanemab LALA-PG (**Fig. 5d-f**). However, no significant increase in IBA1^+^ area around X34^+^ plaques was observed (**Fig. 5g**), suggesting that the rise in OPN reflects an increased expression per cell rather than a higher number of OPN-expressing cells. This observation aligns with our scRNA-seq differential expression analysis (**Fig. 4a**).

### OPN-Driven Aβ Clearance in Lecanemab-Treated Microglia

The proximity of OPN^+^ cells to amyloid plaques suggests that Lecanemab restores protective phagocytic functions in human microglia near Aβ deposits. To test this, we employed our *in vitro* plaque clearance system and stimulated human-derived microglia with increasing concentrations of human OPN (**Fig. 5h-i**). Remarkably, at the highest concentration tested, OPN stimulation significantly decreased the area covered by the pan-Aβ antibody 82E1 (**Fig. 5j**), demonstrating that OPN, one of the main factors induced by Lecanemab, promotes Aβ clearance.

## Discussion

Lecanemab stands as the most successful Aβ-plaque clearing antibody in clinical use^2,13^. Our study demonstrates that its efficacy critically depends on the presence of microglia and the engagement of Fc effector functions, which activate a targeted amyloid-clearing program in these cells. Transcriptomic and functional analyses reveal that Lecanemab induces a distinct program in microglia that enhances phagocytosis and lysosomal activity without triggering the synaptophagy observed in other studies^9^. This selective activation correlates with a reduction in neuritic pathology, potentially underpinning the modest clinical improvements observed with Lecanemab.

Among the Lecanemab-induced upregulated genes, several are highly relevant to AD. For instance, the *MS4A* gene family, including *MS4A6A*, is strongly associated with AD risk, and functions upstream of TREM2^29^. Upregulation of *HSPA5*, an ER stress marker, may indicate enhanced Aβ uptake in Lecanemab-treated microglia, triggering the unfolded protein response^30^. Moreover, *SPP1* stood out as the most strongly upregulated gene in both our scRNA-seq and spatial transcriptomic differential gene expression analyses. However, the overall magnitude of changes in expression levels is relatively limited, which we postulate may be due to the fact that the changes in our microglia are primarily localized to those in close proximity to the plaques. For this reason, despite acknowledging the potential biases of cell-level differential expression analyses, we believe a pseudobulk approach, which has been shown to better control for false discoveries^31^, would mask changes in distinct cell populations.

Nevertheless, despite the subtlety of these changes, our experimental work clearly demonstrates that Lecanemab confers a functional ability to clear plaques. To further extend our investigation of the transcriptional programs associated with this ability, we performed WGCNA, which corroborated our GSEA by revealing that Lecanemab-induced genes cluster into distinct modules associated with the interferon response, antigen presentation, metabolism, and the unfolded protein response. Strikingly, when we leveraged our functional WGCNA modules to analyze spatial transcriptomic changes relative to plaque proximity, we identified the pink module (enriched for *SPP1*) as significantly enriched around Aβ plaques in the Lecanemab-treated samples, further supporting our scRNA sequencing findings. SPP1, a key hub gene in this module, is a well-established marker of disease-associated microglia (DAM/HLA)^16^, cells that accumulate around amyloid plaques without effectively clearing them. Its further induction by Lecanemab suggests the activation of clearance pathways that extend beyond the conventional DAM state. This is supported by our findings that exogenous OPN enhances the amyloid clearance capacity of human microglia. Additionally, we functionally validated one of the top hits identified in both scRNA-seq and spatial GSEA— phagosome/phagocytosis—by demonstrating that Lecanemab promotes Aβ phagocytosis *in vivo* and *in vitro*. Collectively, these findings indicate that a limited set of genes is sufficient to reprogram microglia for efficient amyloid clearance.

Our work highlights the unique advantages of our xenograft human/mouse chimeric model, which allows direct evaluation of the unmodified human antibody on human microglia *in vivo*. Although the Rag2^-/-^ background necessary to prevent graft rejection precludes analysis of adaptive immunity, parallel experiments using a mouse variant of Lecanemab in immune-competent animals confirmed that amyloid clearance is not qualitatively different. This is particularly important given the distinct responses of human versus mouse microglia to amyloid plaques and the low sequence conservation of key AD risk genes like MS4A6A^16^ between species.

A limitation of our study is the limited ability to assess vascular pathology and blood-brain barrier integrity in App knock-in mice models, which have not been reported to exhibit significant defects at this age^32^. However, the observed upregulation in Lecanemab-treated microglia of interferon genes, which have been implicated in brain endothelial dysfunction in AD^33^, suggests a potential mechanism that could underly the amyloid-related imaging abnormalities (ARIA), the most impactful adverse effects of Lecanemab in humans^2^. This also raises the possibility that different Fcγ receptors on microglia (**Extended Data Fig. 4g**, **Table 4**) may differentially mediate the beneficial versus detrimental effects of the antibody. Alternatively, the Fc moiety of Lecanemab may activate Fc receptors or complement receptors in border associated macrophages potentially triggering damaging responses in the vasculature^34^. Future studies will investigate these aspects to better understand the mechanisms underlying risks associated with Lecanemab.

While we cannot rule out the possibility that other anti-Aβ antibodies differentially influence microglia by binding to distinct Aβ species, our findings support the notion that the precise nature of the Aβ protofibrils^1^ targeted by Lecanemab is less critical than its ability to engage amyloid plaques and then microglia through its Fc moiety. This may explain the comparable clinical impact of Donanemab, another FDA-approved antibody that targets a pyroglutamate-modified amyloid peptide in the amyloid plaque itself^3^. Effective amyloid clearance may be achieved if the antibody binds amyloid fibrils sufficiently to correctly position its Fc domain for microglial activation. This insight opens new avenues for therapeutic innovation, including the development of small compounds linked to Fc fragments or the engineering of antibodies with enhanced effector functions and reduced complement activation, an approach extensively utilized in other medical fields^35^, to improve antibody treatment outcomes in Alzheimer’s disease.

## Acknowledgements

This project received funding from the European Research Council (ERC) under the European Union’s Horizon 2020 Research and Innovation Program (grant agreement no. ERC-834682 CELLPHASE_AD). This work was also supported by the Flanders Institute for Biotechnology (VIB vzw), a Methusalem grant from KU Leuven and the Flemish Government (METH/21/05), the Fonds voor Wetenschappelijk Onderzoek (G087523N), the KU Leuven, the Queen Elisabeth Medical Foundation for Neurosciences, the Stichting Alzheimer Onderzoek (SAO) and the Alzheimer’s Association USA (AARF-22-968623). We warmly thank Veronique Hendrickx and Amber Claes for animal husbandry and genotyping; Clare Pridans for providing the construct sequence for generating the Csf1r^ΔFIRE/ΔFIRE^ mice; Hugh Perry for revising the manuscript; Annerieke Sierksma for assisting with statistical analysis; the VIB BioImaging Core where image acquisitions have been performed (in particular, Benjamin Pavie and Nikky Corthout); and Niels Vandamme from the VIB Single Cell Core where sequencing was performed. Graphical schemes were created with BioRender.com. The anti-LAMP1 monoclonal antibody developed by August JT was obtained from the Developmental Studies Hybridoma Bank, created by the NICHD of the NIH and maintained at The University of Iowa, Department of Biology, Iowa City, IA 52242.

## Contributions

G.A. and B.D.S. conceived and designed the study and wrote the manuscript. M.Z. and M.F. conceived and performed bioinformatic analyses. M.D. designed the antibodies sequences, G.A., K.C., M.L.C. and T.J. produced the antibodies in-house. G.A., A.S. and C.X. differentiated human microglial progenitors *in vitro* and performed the *ex vivo* plaque clearance assay; G.A., A.S., C.X. and V.V.L. performed immunofluorescent staining and image acquisition; G.A. and C.X. performed image analysis; L.S. generated the mice; G.A. and A.S. xenotransplanted microglia progenitors into the mouse brain; G.A. and V.V.L. performed antibodies treatment; G.A., K.H., C.X. and A.S. isolated soluble and insoluble brain extracts and performed Aβ MSD; G.A., A.S. and L.W. isolated the human microglia; E.P. supervised the flow cytometry work; L.W., S.P. and M.W. prepared the single-cell libraries; S.P., M.W., G.A. and A.S. performed the Nova-ST experiment with immunofluorescence workflow, K.D. pre-processed the Nova-ST data. All authors discussed the results and commented on the manuscript.

---

**Extended Data Fig. 1.**
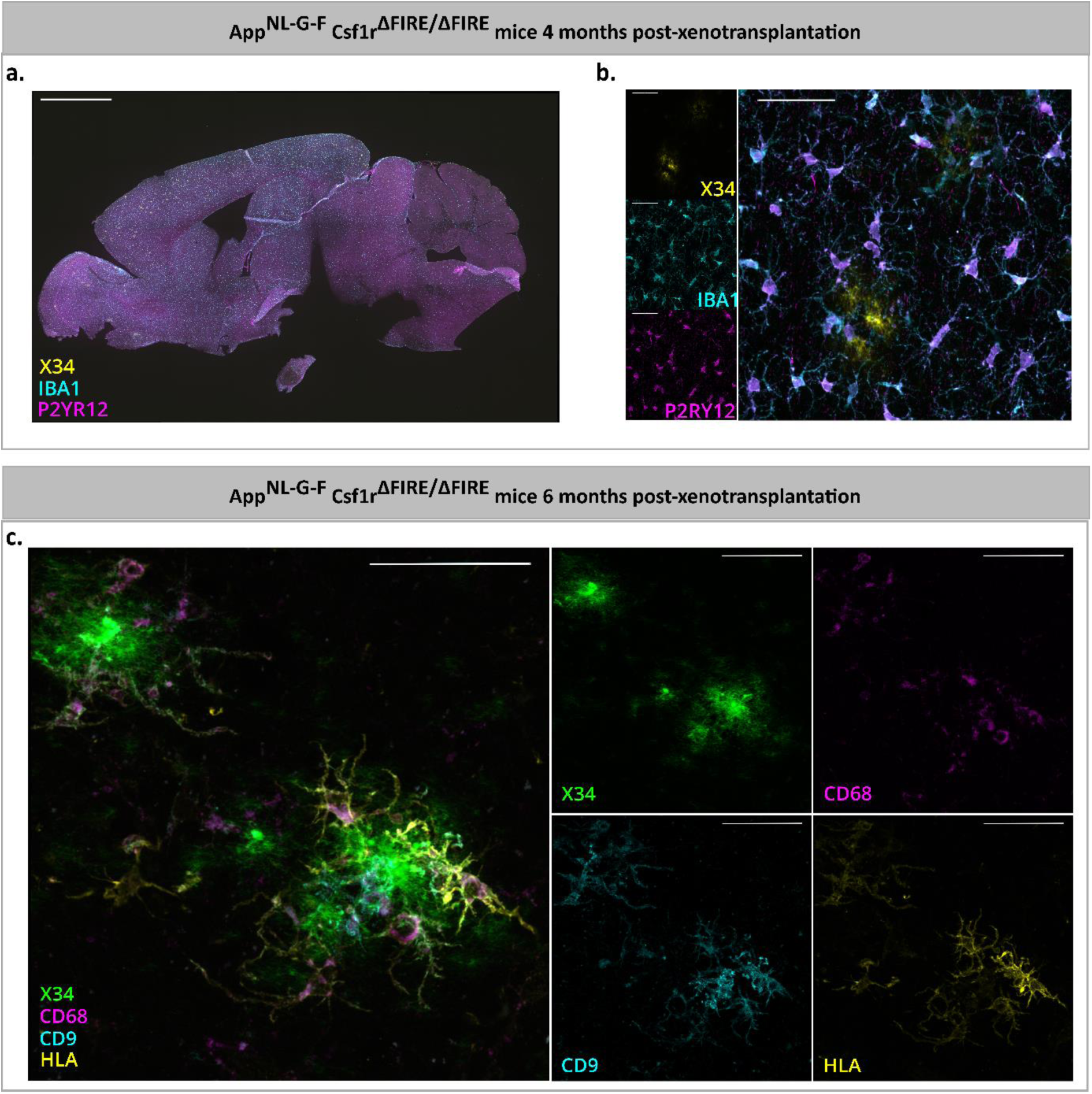
Human microglia in the brain of App^NL-G-F^ mice. **a.** Representative sagittal brain section showing the distribution of human engrafted microglia (labelled with human P2RY12, in magenta, and IBA1, in cyan) in App^NL-G-F^ Csf1r^ΔFIRE/ΔFIRE^ mice at 4 months of age (X34^+^ plaques in yellow). Since these mice genetically lack murine microglia, engraftment efficiency reaches 100%. Scale bar: 1 mm. **b.** Higher magnification confocal image of representative X34 plaques (in yellow) surrounded by human microglia expressing IBA1 (cyan) and P2RY12 (magenta) at 4 months of age. Scale bar: 50 µm. **c.** Representative high magnification confocal z-stacks of X34 plaques (green) surrounded by human microglia expressing CD9 (cyan), HLA (yellow) and CD68 (magenta) at 6 months of age. Scale bar: 50 µm.

**Extended Data Fig. 2.**
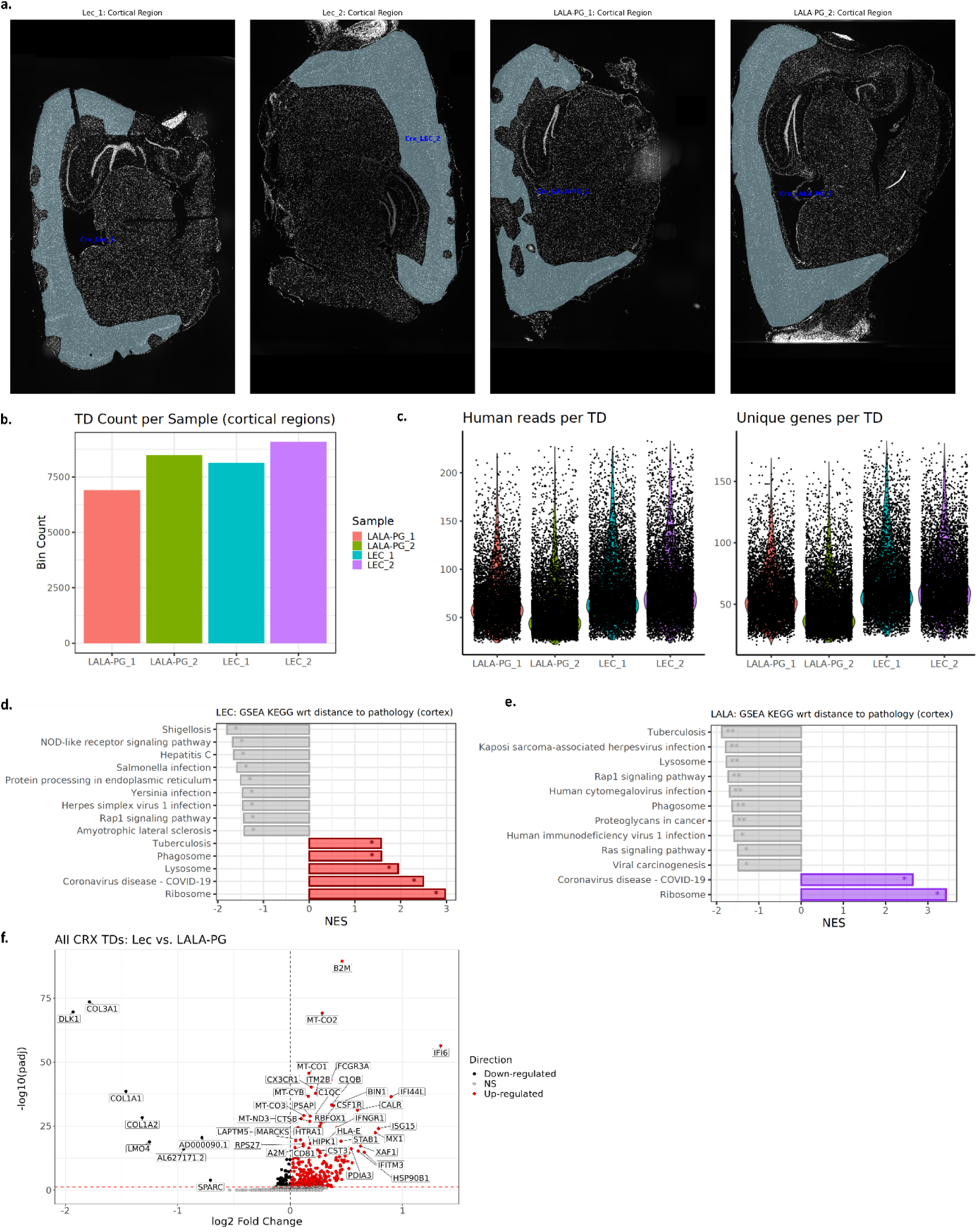
Supplementary information on Nova-ST experiment. **a.** Outline of the cortical regions (blue) selected for subsequent spatial transcriptomic analyses. Regions were manually delineated in QuPath to include cortical areas with high quality stainings in all channels. **b.** Number of tissue domains (TDs, bin40s) per sample, where only TDs falling in specified cortical regions (a) and with at least 30 human transcripts were considered. **c**. Violin plots showing the distribution of number of human UMIs and unique human genes in bin40s across the four samples. **d-e**. Normalized enrichment score (NES) of significantly (p.adj < 0.05) enriched KEGG pathways in function of cortical bins’ proximity to plaques in (**d**) Lecanemab and (**e**) Lecanemab LALA-PG treated mice, as identified by Gene Set Enrichment Analysis (GSEA) on the differential gene expression results in Fig. 1e. (*: p.adj < 0.05, **: p.adj < 0.01, ***: p.adj < 0.001). **f.** Volcano plots showing a gene expression comparison between Lecanemab-and Lecanemab LALA-PG TDs in cortical areas. Adjusted p-value threshold < 0.05 (Wilcoxon rank-sum test, p-values adjusted with Bonferroni correction based on the total number of genes in the dataset, NS = not significant).

**Extended Data Fig. 3.**
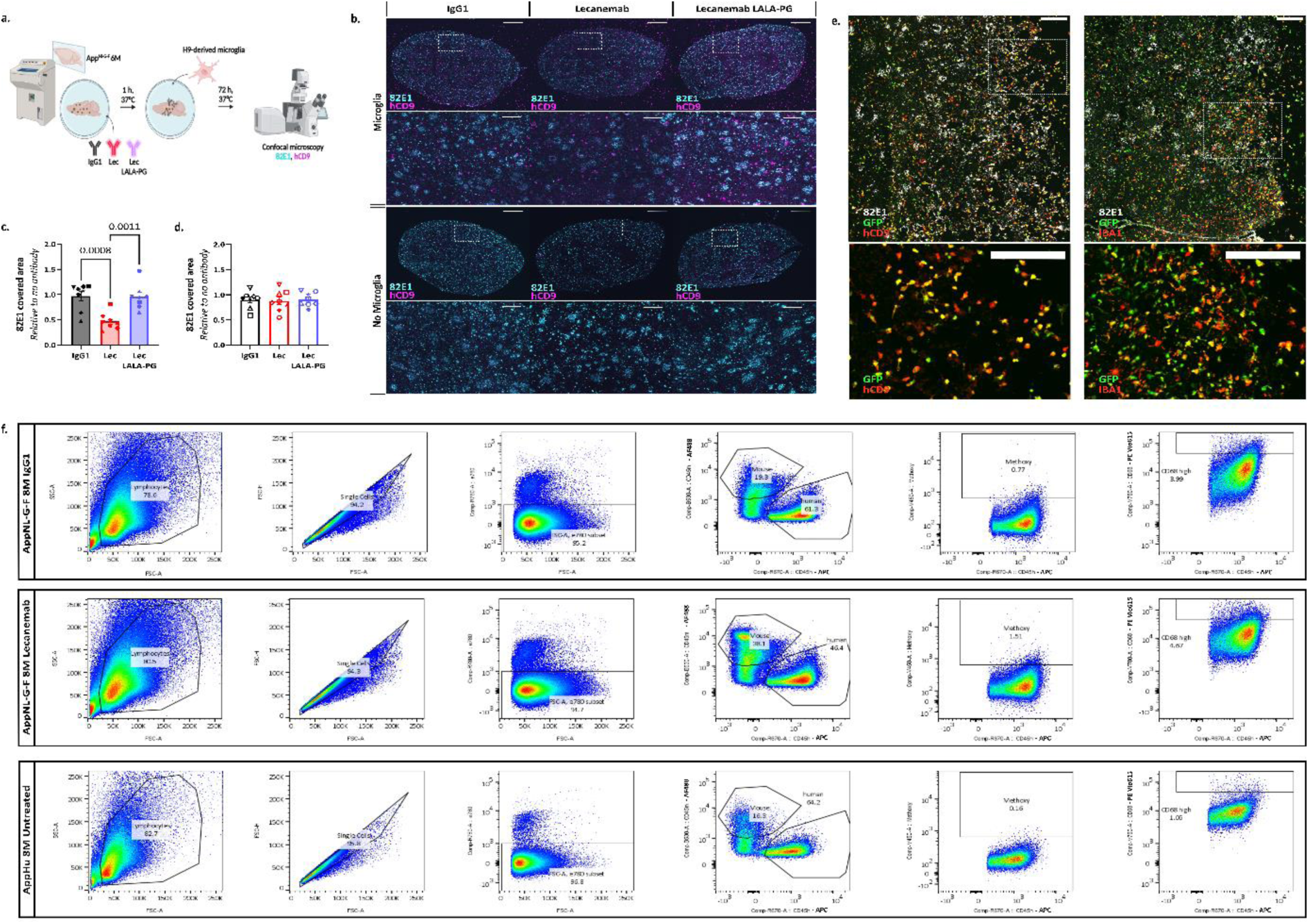
Lecanemab induces plaque clearance in an *in vitro* assay. **a.** Schematic representing the plaque clearance assay paradigm to study Aβ clearance in response to Lecanemab. Sagittal cryosections from App^NL-G-F^ mice at 6 months of age were pre-incubated for 1 hour with IgG1, Lecanemab or Lecanemab LALA-PG. We then added human-derived microglial cells to the cryosections and, after 3 days, quantified the area covered by the pan-Aβ antibody 82E1. **b.** Representative confocal images of 82E1 (Aβ, cyan) and CD9 (human microglia, magenta) immunoreactivity in App^NL-G-F^ brain cryosections after incubation with antibodies as indicated, in presence or absence of human microglia. Scale bar: 1 mm; inset: 200 µm. **c.** Quantification of 82E1 covered area (relative to no antibody condition) in sections plated with human microglia; one-way ANOVA (p = 0.0003). **d.** Quantification of 82E1 covered area (relative to no antibody condition) in sections non plated with human microglia; one-way ANOVA (n.s.). Mean ± SEM shown for each group and points represent independent experiments being each the average of 2-6 cryosections. **e.** Colocalization GFP+ microglia with hCD9 and IBA1. Scale bar: 200 µm. Microglia derived from the H9 stem cell line stably expressing GFP were plated on sagittal cryosections from 6-month-old App^NL-G-F^ mice. After 3 days, cells were fixed to assess GFP^+^ expression of hCD9 and IBA1. The majority of GFP^+^ cells co-express hCD9 (left panels), while not all GFP^+^ cells are IBA1^+^ (right panels). **f.** Representative FACS plots of methoxy-X04⁺ human microglia from 8-month-old App^NL-G-F^ mice injected with IgG1 (top panels) or Lecanemab (middle panels), along with untreated App^Hu^ mice as a negative control (bottom panels). Live human cells were identified based on CD11b⁺ hCD45⁺ expression, with the negative control used to establish the gating for methoxy-X04⁻ cells. Methoxy-X04⁺ cells were quantified as a percentage of hCD45⁺ human microglia and within the CD68^high⁺ population (Fig. 3a**-c**).

**Extended Data Figure 4.**
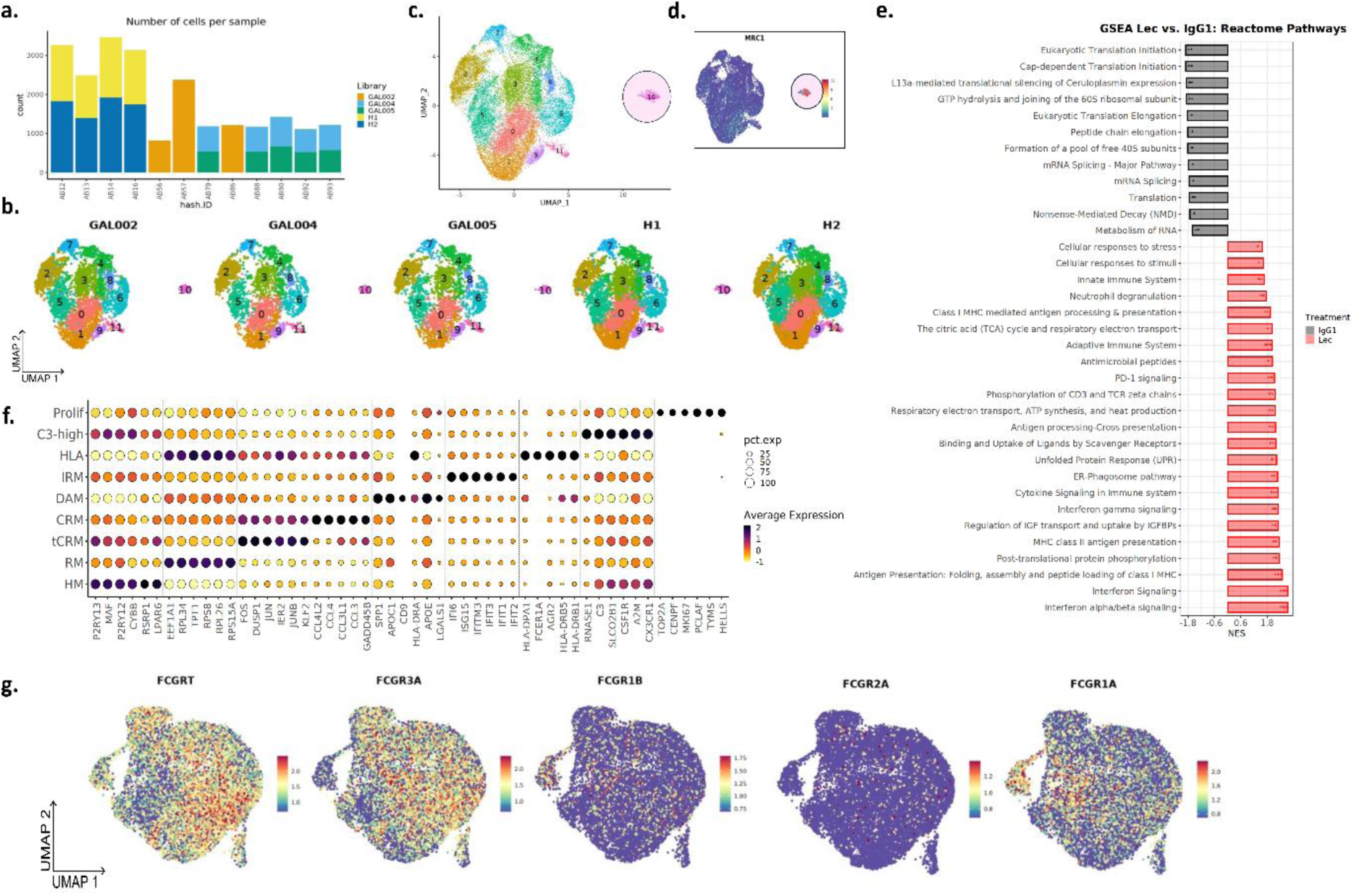
Preparation of the datasets for analysis and cell type/state annotations. **a-b.** Overview of the cDNA libraries used in single-cell RNA-seq experiments (GAL002, GAL004, GAL005, H1 and H2), including the number of cells per sample (**a**) and the distribution of cells on the UMAP across different libraries after data integration (**b**). **c.** The UMAP plot of the 22,841 single xenografted human cells that passed quality control colored by clusters. **d.** UMAP plot colored by the expression level of MRC1, a macrophage marker. Notably, Cluster 10, identified as the macrophage cluster, was removed from further analysis. **e.** Reactome pathway enrichment terms for significantly downregulated (black) and upregulated genes (red) ranked by p-value upon Lecanemab treatment. **f.** Top six most differentially expressed genes in each cell state. The dot size represents the percentage of cells within the given cluster expressing the gene. The dot color represents the scaled and normalized average expression of the gene. **g.** UMAP plots colored by the level of expression of *FCGRs* (encoding FcγRs) expressed by human microglia.

**Extended Data Figure 5.**
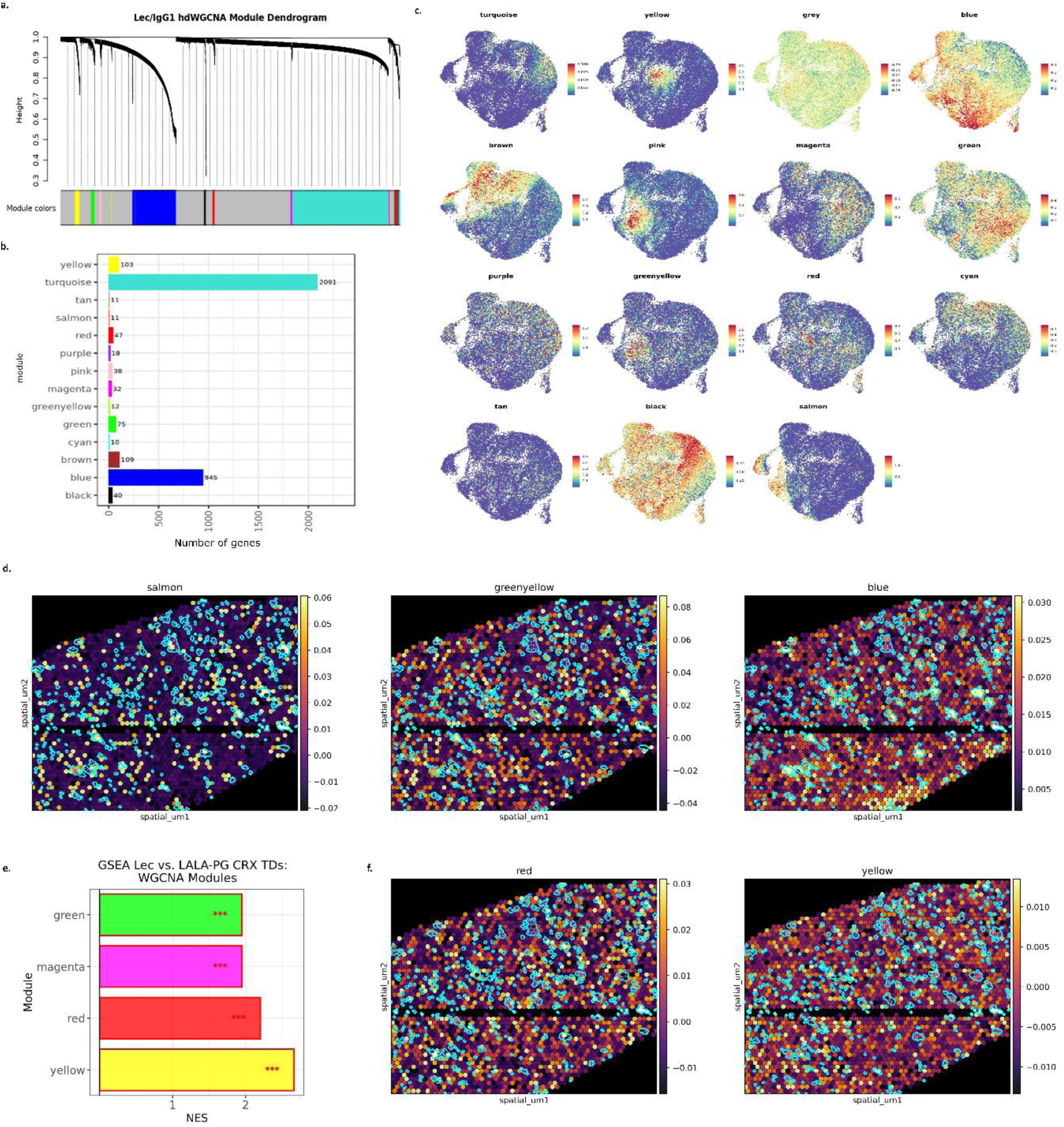
Lecanemab affects pathways linked to unfolded protein response, cell metabolism, antigen presentation and interferon response. **a.** Weighted Gene Co-expression Network Analysis (WGCNA) cluster dendrogram groups genes measured across IgG1- and Lecanemab-treated cells into 14 distinct modules, defined by dendrogram branch cutting. An additional "unassigned" gray module was identified but discarded from subsequent analyses. **b.** Number of genes belonging to identified WGCNA modules. **c.** UMAP plots colored by the combined level of expression of each WGCNA module. **d.** Cortical tissue domains (TDs) in Lecanemab-treated mice colored based on their relative expression of genes belonging to the blue (left panel), salmon (middle panel) and greenyellow (right panel) modules (purple: low expression, yellow: high expression) and overlayed with plaque ROIs (outlined in cyan). Enrichment scores were obtained using scanpy’s *score_genes()* function. Note the significant enrichment of the module in bins close to Aβ plaques, as quantified in Fig. 2b**. e.** NES of significantly enriched (p.adj < 0.05) Weighted Gene Co-expression Network Analysis (WGCNA) modules between Lecanemab and Lecanemab LALA-PG treated TDs in cortical regions, as identified by GSEA (***: p.adj < 0.001). **f.** Cortical TDs in Lecanemab-treated mice colored based on their relative expression of genes belonging to the red (left) and yellow (right panel) modules (purple: low expression, yellow: high expression) and overlayed with plaque ROIs (outlined in cyan).

**Table 1 Human microglial transcriptomic modules identified by Weighted Correlation Network Analysis (WGCNA).**

**Table 2 Gene Ontology (GO) analyses of the human microglial transcriptomic modules identified by Weighted Correlation Network Analysis (WGCNA) (top 20 terms).**

**Table 3 Overlap between the human microglial transcriptomic modules identified by Weighted Correlation Network Analysis (WGCNA) and the ATM**^26^ **and PAM**^27^.

**Table 4 Differential expression of FCGRs in the human microglia.**

