## Supplementary material for "The Alzheimer’s therapeutic Lecanemab induces an amyloid-clearing program in microglia": Materials and Methods

#### Design and preparation of monoclonal anti-Aβ antibodies

For the experiments described below, Lecanemab, Lecanemab engineered to lack effector function (Lecanemab LALA-PG) or control human IgG1 were used. First, the amino acid sequences for the variable domains of Lecanemab were retrieved from the KEGG DRUG Database (**Table S1**). Heavy and light chain were cloned into a single IgG1 expression vector to express a fully human IgG1 antibody. For Lecanemab, the production was initially done in-house, then outsourced to Genscript. Briefly, synthetic genes encoding for the respective variable domains, preceded by the mouse Ig heavy leader signal (Twist Biosciences) were cloned in pTRIOZ-hIgG (Invivogen). The VL domain was cloned by using restriction enzymes AscI/BsiWI, the VH domain by AgeI/NheI. The antibody encoding open reading frames were sequence confirmed by Sanger sequencing (Eurofins). Plasmid DNA was delivered to CHO cells (Thermofisher, #A29127) by transient transfection according to manufacturer's protocol (CHOgro High Yield Expression System, Mirus Bio, #MIR 6270). Transfected CHO cells were cultured for 14 days in suspension on agitation at 32°C. Two weeks post-transfection, cell supernatants were collected and incubated overnight at 4°C with AmMag™ Protein A Magnetic Beads (GenScript, #L00939). The beads were collected using a magnetic separation rack and targeted antibodies were separated using the AmMag™ SA Plus system (GenScript,

#L01013). To abolish Fc effector function, the heavy chain of Lecanemab was designed to include the P329G substitution combined with L234A and L235A (LALA-PG). The light chain was the same as for Lecanemab (**Table S1**). Production was outsourced to Genscript. As control, we used the human IgG1 isotype control (Imtec Diagnostics, #LT9005). For the murine antibodies (mAb158 and mAb158 LALA-PG), production was outsourced to Genscript. The amino acid sequence mAb158 was retrieved from patent US 8025878 B2, seq IDs 115 and 116. The LALA-PG mutations were designed according to Schlothauer et al.[1] As control, we used the mouse IgG2a isotype control (Leinco, #P381). The purity of the antibodies was estimated to be above 75% by densitometric analysis of the Coomassie Blue-stained SDS-PAGE gel under non-reducing conditions. Binding to Aβ1-42 (rPeptide, #A-1163-2), was confirmed by ELISA and Dot Blot as performed in[2,3].

**Human microglial progenitor differentiation and xenotransplantation into the mouse brain**

Human embryonic stem cells (WiCell Research Institute, RRID: CVCL_9773, #WA09) were differentiated into microglial precursors as previously described[4]. Routine culturing and maintenance of the stem cells was performed on Matrigel coated surface (Corningusing E8-flex growth media (Thermo, #A2858501). Cells were maintained in a humidified chamber at 37°C with 5% CO2. Once cells reached approximately 70–80% confluence, stem cell colonies were dissociated into single cells using Accutase (Sigma-Aldrich) and plated into U-bottom 96-well plates at a density of about 10,000 cells per well in mTeSR1 medium supplemented with BMP4 (50 ng $ml^{-1}$), VEGF (50 ng $ml^{-1}$), and SCF (20 ng $ml^{-1}$) for 4 days, allowing them to self-aggregate into embryoid bodies. On day 4, embryoid bodies were transferred into six-well plates (approximately 20 embryoid bodies per well) in X-VIVO (Lonza, #LO02-060F,) medium with supplements, and further supplemented with SCF (50 ng $ml^{-1}$), M-CSF (50 ng $ml^{-1}$), IL-3 (50 ng $ml^{-1}$), FLT3 (50 ng $ml^{-1}$), and TPO (5 ng $ml^{-1}$) for 7 days. A full medium change was performed on day 8. On day 11, the differentiation medium was replaced with X-VIVO (plus supplements) containing FLT3 (50 ng $ml^{-1}$), M-CSF (50 ng $ml^{-1}$), and GM-CSF (25 ng $ml^{-1}$). On day 18, floating human microglial precursors were collected from the supernatant and resuspended in Dulbecco's Phosphate Buffered Saline (PBS, Gibco) at a concentration of 2.5 x $10^5$ cells/µL and engrafted into P4 mouse brains (5 x $10^5$ cells per pup) as previously described[5]. Embryoid bodies were re-transferred into six-well plates in X-VIVO (plus supplements) containing FLT3 (50 ng $ml^{-1}$), M-CSF (50 ng $ml^{-1}$), and GM-CSF (25 ng $ml^{-1}$) for further collections. All cytokines were purchased at Pepro Tech.

**Mice and antibody treatment**

Homozygous mouse oocytes from $Rag2^{tm1.1Flv}$; $Csf1^{tm1(CSF1)Flv}$; $Il2rg^{tm1.1Flv}$; $App^{tm3.1Tcs}$ crosses were microinjected with reagents targeting the fms-intronic regulatory sequence (FIRE sequence) in the intron 2 of the mouse Csf1R gene (PMID: 31324781). Ribonucleoproteins containing 0.3 µM purified Cas9HiFi protein 0.3 µM crRNA ( 5'GTCCCTCAGTGTGTGAGA3' and 5'CAATGAGTCTGTACTGGAGC3') and 0.3 µM trans-activating crRNA (Integrated DNA Technologies) were injected into the pronucleus of 120 embryos by the CBD Mouse Expertise Unit of KU Leuven. One female founder with the expected 428 bp deletion was selected and crossed with a $Rag2^{tm1.1Flv}$; $Csf1^{tm1(CSF1)Flv}$; $Il2rg^{tm1.1Flv}$; $App^{tm3.1Tcs}$ male and the progeny were interbred to obtain a $Rag2^{tm1.1Flv}$; $Csf1^{tm1(CSF1)Flv}$; $Il2rg^{tm1.1Flv}$; $App^{tm3.1Tcs}$; $Csf1R^{em1Bdes}$, named hereafter as $App^{NL\text{-}G\text{-}F}$ $Csf1r^{\Delta FIRE/\Delta FIRE}$. For maintaining the colony, $App^{NL\text{-}G\text{-}F}$ $Csf1r^{\Delta FIRE/\Delta FIRE}$ males were crossed with App$^{NL\text{-}G\text{-}F}$ Csf1r$^{\Delta FIRE/WT}$ females as 5 times homozygous females tend to take less care of their progeny. For grafting, App$^{NL\text{-}G\text{-}F}$ Csf1r$^{\Delta FIRE/\Delta FIRE}$ pups were fostered to CD1 mothers to facilitate their survival. App$^{NL\text{-}G\text{-}F}$ mice (App$^{tm3.1Tcs+}$; Takaomi Saido[6]) express amyloid precursor proteins (APP) at endogenous levels but contain the humanized Aβ sequence along with Swedish (NL; K670_M671delinsNL), Arctic (G; E693G), and Iberian (F; I716F) familial Alzheimer's disease-causing mutations in C57BL/6 background. A cohort of Rag2$^{tm1.1Flv}$; Csf1$^{tm1(CSF1)Flv}$; Il2rg$^{tm1.1Flv}$; App$^{tm3.1Tcs}$ mice (App$^{NL\text{-}G\text{-}F}$ mice) was xenotransplanted with human microglia[5], treated between 6 and 8 months of age, and used for the *in vivo* phagocytosis assay. To establish proper gating for this experiment, we also included also Rag2$^{tm1.1Flv}$; Csf1$^{tm1(CSF1)Flv}$; Il2rg$^{tm1.1Flv}$; App$^{em1Bdes}$ mice (App$^{Hu}$ mice) as negative controls. These mice have humanized Aβ sequence[7] and do not develop Aβ pathology. Like App$^{NL\text{-}G\text{-}F}$ mice, they were xenotransplanted with human microglia. Since neither App$^{Hu}$ nor App$^{NL\text{-}G\text{-}F}$ mice are genetically devoid of endogenous mouse microglia, these cells were depleted before transplantation by inhibiting the CSF1 receptor (CSF1R) using BLZ945 at a dose of 200 mg/kg on postnatal days 2 and 3 (P2 and P3), as previously described[5]. Mice had access to food and water ad libitum and were housed with a 14/10 h light-dark cycle at 21°C in groups of two to five animals. All experiments were conducted according to protocols approved by the local Ethical Committee of Laboratory Animals of the KU Leuven (government license LA1210579, ECD project numbers P125/2022 and P132/2022) following country and European Union guidelines.

The ability of Lecanemab (or mAb158) to reduce Aβ burden was assessed following chronic treatment of 4-month-old male and female App$^{NL\text{-}G\text{-}F}$ Csf1r$^{\Delta FIRE/\Delta FIRE}$ mice (or App$^{NL\text{-}G\text{-}F}$ mice) dosed weekly intraperitoneally for 8 weeks with 10 mg $kg^{-1}$ Lecanemab (mAb158), Lecanemab LALA-PG (mAb158 LALA-PG) or IgG1 (IgG2a). One day after the final dose, mice were terminally anesthetized with an overdose of sodium pentobarbital and transcardially perfused with ice-cold 1 × DPBS (Gibco, #14190-144) supplemented with 5 U of heparin (LEO). The brain was quickly removed and halved along the medio-sagittal line. The right hemisphere was fixed by immersion in formaldehyde solution 4% for 24 h, followed by storage in PBS solution with 0.01% sodium azide until being cut. The left hemisphere was or i) snap-frozen for MSD ELISA, or ii) immediately processed for human microglia isolation. The mice used for analysing antibodies distribution and Nova-ST experiment were transcardially perfused with ice-cold PBS supplemented with 5 U of heparin; then i) one hemisphere was snap-frozen for MSD ELISA and ii) the other was embedded in cold OCT separately and snap-frozen in isopentane chilled with liquid nitrogen. Samples were stored at -80°C.

### Spatial Transcriptomics with Nova-ST

#### Tissue collection for Nova-ST with immunofluorescence

Tissue collection was performed as described in[8]. Briefly, OCT-embedded hemispheres were cryosectioned sagittaly using a CryoStar NX70 cryostat (ThermoFisher). First, 5–10 serial sections of 50 µm thickness were cryo-sectioned from a region farther away from the region of interest. These serial sectioned OCT scrolls were put into a 2 mL lo-bind tube (Eppendorf Cat. No. 0030108078) and stored at -80°C. The tissue scrolls were washed with 1 mL of ice-cold PBS at 4°C. Total RNA extraction from the spun-out tissue was performed using innuPREP mini-RNA kit (Analytik Jen; Cat. No. AJ 845-KS-2040250). Manufacturer recommendations were followed to extract total RNA. Elution was performed in 30 mL of nuclease free water, NFW (Thermo Fisher; Cat No. 10977035). The quality of the total RNA was assessed using Pico RNA kit (Agilent) and was above 8 for all samples.

**Optimization of tissue permeabilization for Nova-ST with immunofluorescence**

The tissue optimization for Nova-ST is performed using 10X Genomics Visium Spatial Tissue Optimization kit, according to the manufacturer's protocol (Visium Spatial Tissue Optimization Reagents kit, User Guide CG000238.Ref F). Briefly, 8x 10 µm tissue sections are placed on the 8x available slots on the Visium Spatial Tissue Optimization Slide (PN: 300394). Methanol fixation, immunofluorescence staining and Imaging was performed as described in the 10X Genomics demonstrated protocol (CG000312.RevF). The permeabilization test is performed as per manufacturer's recommendations, except the permeabilization enzyme (PN: 2000214) was replaced by 0.65U/µl working stock of Pepsin enzyme in 0.01N HCl (pH 2.0) (Sigma Aldrich; P7000-25G). The optimal permeabilization time is determined by performing the permeabilization time series of eight different time points ranging from 3 mins – 45 mins. The optimal permeabilization time thus determined is used for the downstream spatial transcriptomics workflow[9] (further details in[8]). Similar to other spatial techniques such as 10X Genomics Visium & BGI STomics, the optimal permeabilization time is assessed based on the strongest fluorescence signal with the lowest signal diffusion (crispness of the RNA footprint). Based on our assessment, we found the most optimal permeabilization time for the mouse brain section to be 14 mins.

**Spatial Transcriptomics analysis using Nova-ST**

The Nova-ST spatial transcriptomics analysis is performed using Nova-ST chips prepared from repurposed NovaSeq 6000 S4 flow cell as described in the publication[9] and resources[10]. Spatial transcriptomics workflow is followed as per the publication[9] (additional details in[8]). Briefly, Nova-ST chip was removed from the storage buffer and washed with nuclease free water and dried at 37°C. Next, a 10 µm tissue section from the desired region of interest (lateral 1.56 mm) was prepared from the tissue cryo-block and placed on the Nova-ST chip and thawed to attach the tissue layer on the surface of the chip. The chip was then stored at -80°C until the spatial transcriptomics experiment. For performing the ST assay, the tissue was dried on a 37°C hot plate for 2 mins and the chip was then dipped into 100% methanol at -20°C and incubated for 30 mins to fix and permeabilize the tissue. After the methanol fixation, the Nova-ST chip was blocked using 200 µl of blocking buffer (2X blocking buffer prepared with 6X SSC buffer, Sigma Aldrich #S6639-1L; 4% BSA, Sigma Aldrich #A9576-50ML; 0.2% TX-100, Sigma Aldrich #X-100-5ML and 10% (v/v) Ribonucleoside Vanadyl Complex, NEB # S1402S) for 5 minutes. After removing the blocking buffer, we added 110 µl of conjugated anti-IgG primary antibody staining mix (2X blocking buffer; goat anti-Human IgG- Alexa Fluor™ 647: 80 $\mu g\,ml^{-1}$, #A-21445, ThermoFisher Scientific; 6U/ µl of RNAse inhibitor, NEB #M0314L) and is incubated for 30 mins. After the incubation, the chip was washed 5x times with wash buffer (2X blocking buffer & 10% (v/v) Ribonucleoside Vanadyl Complex, NEB # S1402S). After the last wash with the buffer, we added 110 µl of conjugated anti-hCD45 and anti-D54D2 antibodies staining mix (2X blocking buffer; rabbit anti-β-Amyloid (D54D2)- Alexa Fluor™ 594: 80 µg $ml^{-1}$, #35363S, Bioké; mouse anti-Human CD45- Alexa Fluor™ 488: 80 µg $ml^{-1}$, #304017, BioLegend; 80 µg/mL, 6U/ µl of RNAse inhibitor, NEB #M0314L) and incubated for 30 mins. After the incubation, the chip was washed 5x times with wash buffer (2X blocking buffer & 10% (v/v) Ribonucleoside Vanadyl Complex, NEB # S1402S). The chip was then wash by dipping 20X times in 3X SSC buffer. 20 µl of glycerol mounting medium (85% glycerol, Sigma Aldrich # G5516-100ML and 2U/µl of RNAse inhibitor, NEB #M0314L) was added onto the tissue on the chip and after placing a coverslip, the tissue on the Nova-ST chip was imaged for the DAPI and antibody stains. For downstream tissue alignment and analysis of the spatial transcriptomics data , images were registered for using Nikon NiE 8-staged driven by NIS-Elements AR (6.10.01) software, using a 10x/0.45 air objective (Plan Apo 10x Lambda CFI Eclipse DIC N1, Nikon #MRD00100). Images had a resolution of 2048 x 2048, and 0.63 µm pixel size. Additional high magnification images were taken with Nikon AX Confocal Microscope System driven by NIS-Elements AR (5.41.01) software, using a 40x/0.75 air objective (Plan Apo Lambda S 40XC Sil, Nikon # MRD73400). Ten stacks were acquired using a 0.8 µm z-interval and images had a resolution of 2048 x 2048, and 0.12 µm pixel size. 3D surface reconstructions were performed with Imaris Software (Bitplane).

After imaging, the coverslip was removed from the chip and the chip washed with excess of 3X SSC buffer. After incubating the chip in 3X SSC for 3 minutes, the chip was subject to tissue permeabilization with 120 µl of tissue permeabilization mix of 0.65U/µl working stock of Pepsin enzyme in 0.1N HCl (pH 2.0) (Sigma Aldrich, #P7000-25G) at 37°C. The optimal permeabilization time estimated from the tissue permeabilization analysis (14 mins) was used for the transcriptomics analysis. After washing the chip with successive rounds of 0.1X SSC wash buffer and 1X RT wash buffer, the reverse transcription mix (1.05 U/µl RNAse Inhibitor, Lucigen, #30281-1; 4.5% Ficoll PM-400, Sigma, #F5415-50 ML; 1.05 mM dNTPs 25 mM, Thermo Fisher, #R1121; 10 U/µl Maxima h-RT, Thermo Fisher, #EP0753) was added to the chip and incubated at 42°C overnight (18-24 hrs) to complete the reverse transcription. The chip was then subjected to successive rounds of wash with 0.1X SSC and 1X Exo I buffer. Chip was then subjected to exonuclease digestion (Exonuclease I, NEB, #M0293L) at 37°C for 45 minutes followed by tissue removal digestion (100 mM Tris pH 8.0, Thermo Fisher, #J22638.AE; 200 mM NaCl, Thermo Fisher, #AM9760G; 2% SDS, Thermo Fisher, #AM9822; 5mM EDTA, Thermo Fisher, #AM9260G and 16 mU/µl of Preteinase K, NEB, #P8107S) to remove the leftover tissue from the Nova-ST chip by incubating the chip in the tissue removal mix at 37°c for 60 minutes. Caustic denaturation (0.1 N NaOH pH 13, Sigma Aldrich, #72068-100ML) was performed on the chip to remove the RNA strands from the first strand product on the Nova-ST chip. After the 3 rounds of neutralization washes (100 mM Tris pH 7.5, Thermo Fisher, #15567027) and nuclease free water washes, the chip was then subjected to second strand synthesis reaction. In this reaction the chip was submerged into second strand synthesis mix (1X NEB buffer 2, NEB #M0212L; 11 uM RPE randomer, IDT technologies; 0.55U/µl of Klenow exo (-) fragment, NEB M0212L) and incubated for 120 mins. The second strand cDNA product was extracted from the Nova-ST chip using caustic denaturation (0.1 N NaOH pH 13, Sigma Aldrich #72068-100ML). The denatured product was neutralized with 100 mM Tris pH 7.0, (Thermo

Fisher, #AM9850G) and subsequently purified with 1.8X Ampure XP beads (Beckman Coulter, #A63882) using the manufacturer's recommendation. The purified library was PCR amplified and purified as described in the protocol. After the quality assessment using bioanalyzer (Agilent) and Qubit (Thermo Fisher) library normalization was performed. Sequencing library preparation was performed using indexed PCR amplification of the library. Library purification was performed with 2X successive rounds of Ampure XP purification (0.8X followed by 1.0X purification). The rest of purification protocol was performed as per manufacturer’s recommendations.

Nova-ST libraries were sequenced with NextSeq 2000 (Illumina) for shallow sequencing (for quality estimation of the sample) and MGISEQ-2000 (BGI) sequencing platforms for deeper sequencing. Sequencing of the Nova-ST libraries were performed with paired end sequencing of read of 34 bps (read 1), 85 bps (read 2), 8 bps (index 1) and 8 bps (index 2). MGISEQ-2000 sequencing was done at the MGI Hong Kong SAR sequencing facility. Before being able to sequence on the MGI platforms, the Nova-ST libraries needed to undergo a conversion step using MGIEasy Universal DNA Library Prep kit. Briefly, the sequencing libraires were circularized using the splint-ligation step and the circularized libraries were converted to single stranded DNA copies. DNA nanoballs were prepared from the circularized ssDNA using Rolling Circle Amplification (RCA). DNB libraries generated were then flown through the patterned flow cell of the MGISEQ-2000RS High-Throughput Sequencing kit. For a targeted sequencing saturation of 50-60%, sequencing were performed to a depth of 1 billion reads per sample.

**Region of interest detection**

Single fluorescent and brightfield images were stitched using Fiji (MultiView-Stitcher) with a 15% overlap. The resulting overlays of immunostainings and deep-sequencing images were imported into QuPath to identify cortical regions. Within these regions, we selected regions of interest ($D54D2^{+}$ plaque regions) based on a size exclusion criterion of $<20\ \mu m^2$. A semi-automated script in QuPath (v0.4.3) was used to detect plaque ROIs, which were then manually screened to exclude artifacts. Finally, GEOjson files were exported for subsequent analysis.

**Nova-ST data alignment, quality control, and pre-processing**

A combined Human (hg38) and Mouse (mm10) STAR index was created from the 10x Genomics GRCh38-and-mm10-2020-A index using STAR (v2.7.11a). This index was used together with the NovaScope Pipeline v1.1[11], utilizing spatula (v1.0.0) and STAR (v2.7.11a) to pre-process, map and quantify the data via the following two workflows: sge-per-run, transcript-per-unit. Mapped and quantified data was loaded and binned into hexagonal bins (TDs) with a width, and center-to-center distance of 40µm using custom Python code, images were exported at a resolution of 1px:1µm and used for alignment with immunofluorescent images previously collected. Images were aligned to spatial data using the Landmark Correspondences plugin in Fiji. Multiple fiducial circles, visible in both the 488 IF channel and the spatial data, were selected as landmarks. An affine transformation was then applied to achieve precise alignment.

Expression matrices of all TDs were read into Scanpy[12] and pre-processed. Genes were subset to retain only those mapping to the human transcriptome. Quality control metrics were calculated and TDs with fewer than 30 UMIs and greater than 250 UMIs were removed. GEOjson files specifying cortical regions and plaque ROIs were read into python using geopandas and overlapped with the spatial object (Extended Data Fig. 2a). TDs falling in the cortical region were retained and the distance from the center of each TD to the edge of the nearest segmented plaque was calculated using Scipy's KDTree package: (https://github.com/scipy/scipy/blob/main/scipy/spatial/kdtree.py).

**Distance-based differential expression analysis (DE)**

DE analyses of TDs with respect to distance to pathology were conducted by fitting generalized linear models (GLM) in the Lecanemab and Lecanemab LALA-PG treated mice separately. Analysis was performed only on TDs within 200µm of a plaque edge to limit the biasing effect of outlying TDs, and on genes expressed in at least 90 TDs. Each GLM model was tested for DE with EdgeR's (v3.40.0) quasi-likelihood F-test (QLFTest) which accounts for the uncertainty in dispersion estimation. Multiplicity correction was performed by applying the Benjamini-Hochberg (BH) method on the associated p-values, and a significance threshold of p.adj <0.05 was used for all DEs. Gene Set Enrichment Analyses (GSEAs) were performed as specified in the "Single-cell sequencing data analysis" section.

**Visualization of gene set expression levels in TDs**

To visualize the expression level of gene sets in TDs, TD expression levels were first normalized using Scanpy's *sc.pp.normalize_total()* and *sc.pp.log1p()* functions. Relative expression levels of gene sets were determined using the *sc.tl.score_genes()* function, and *sc.pl.spatial()* was used to spatially visualize the resultant scores overlaid over the plaque segmentation image (Fig. 5c, Extended Data Figs. 5d,f).

**Immunofluorescence on vibratome sections**

Thirty µm-thick sagittal slices were cut using a vibratome (Leica VT1000S). At least 2 brain slices per mouse covering the regions of interest (lateral 1.56 mm) were selected following the mouse brain atlas of Franklin and Paxinos[13]. After 3 washes in PBS, antigen retrieval was performed by microwave boiling in 10 mM tri-sodium citrate buffer pH 6.0. Slices were then permeabilized in PBS containing 0.2% Triton X-100 for 15 minutes. After permeabilization, slices were stained with X-34 staining solution (10 µM X-34 (Sigma-Aldrich), 20 mM NaOH (Sigma-Aldrich), and 40% ethanol in PBS) for 20 min at room temperature. Slices were washed three times with 40% ethanol in PBS for 2 min and twice with PBS + 0.2% Triton for 5 min. Afterwards, slices were blocked with PBS-T with 5% donkey serum at room temperature for 1h and stained with primary antibodies overnight at 4 °C with gentle agitation (for synaptic staining, anti-Synaptophysin and anti-Homer 1 primary antibodies were incubated for 2 consecutive nights). Primary antibodies, rabbit anti-human P2YR12 (0.2 µg $ml^{-1}$; Atlas Antibodies, #HPA013796), mouse anti-amyloid beta (N) (clone 82E1) antibody (0.2 µg $ml^{-1}$; IBL, #10323), guinea pig anti-IBA1 antibody (2 µg $ml^{-1}$; Synaptic systems, #234308), rat anti-LAMP1 (4 µg $ml^{-1}$; DSHB, #1D4B-c), rabbit anti-IBA1 antibody (1 µg $ml^{-1}$; Wako, #019-19741), rabbit anti-Homer 1 antibody (2 µg $ml^{-1}$; Synaptic systems, #160003), mouse anti-

Synaptophysin antibody (2 µg ml$^{-1}$; Synaptic systems, #101011), goat anti-osteopontin/OPN (2 µg ml$^{-1}$; R&D System, AF808), were diluted in PBS-T with 5% donkey serum. The next day, after rinsing three times with PBS-T, secondary fluorochrome-conjugated antibodies were added to PBS-T with 5% donkey serum for 2 h at room temperature. The following secondary antibodies were used: goat anti-rabbit, Alexa Fluor™ 647 (2 µg ml$^{-1}$; Invitrogen, #A21245); goat anti-mouse, Alexa Fluor™ 647 (2 µg ml$^{-1}$; Invitrogen, #A21236); goat anti-guinea pig, Alexa Fluor™ 488 (2 µg ml$^{-1}$; Jackson Immunolab, #106-545-003); goat anti-rat, Alexa Fluor™ 594 (2 µg ml$^{-1}$; Invitrogen, #A11007); donkey anti-mouse, Alexa Fluor™ 647 (2 µg ml$^{-1}$; Invitrogen, #A31571); donkey anti-rabbit, Alexa Fluor™ 594 (2 µg ml$^{-1}$; Invitrogen, #A21207); goat anti-mouse, Alexa Fluor™ 546 (2 µg ml$^{-1}$; Invitrogen, A21123); donkey anti-goat, Alexa Fluor™ 488 (2 µg ml$^{-1}$; Invitrogen, A11055). After rinsing three times with PBS-T and once with PBS, sections were mounted onto glass slides using the Glycergel mounting media and allowed to dry at room temperature.

For showing the colocalization of HLA, CD9 and CD68, after X34 staining, wash and blocking, slices were incubated overnight at 4 °C with gentle agitation with mouse anti-human HLA antibody (5 µg ml$^{-1}$; Abcam, ab7856). The next day, after rinsing three times with PBS-T, goat anti-mouse, Alexa Fluor™ 647 (2 µg ml$^{-1}$; Invitrogen, #A21236) was added to PBS-T with 5% donkey serum for 2 h at room temperature. Sections were washed, blocked and incubated with the following primaries overnight at 4 °C with gentle agitation: mouse anti-human CD9 biotinilated (10 µg ml$^{-1}$; Biolegend, #312112) and rabbit anti-human CD68 (4 µg ml$^{-1}$; Abcam, #ab213363). The next day, after rinsing three times with PBS-T, secondary fluorochrome-conjugated antibodies were added to PBS-T with 5% donkey serum for 2 h at room temperature. The following secondary antibodies were used: donkey anti-rabbit, Alexa Fluor™ 594 (2 µg ml$^{-1}$; Invitrogen, #A21207); streptavidin, Alexa Fluor™ 488 (2 µg ml$^{-1}$; Invitrogen, A11055). After rinsing three times with PBS-T and once with PBS, sections were mounted onto glass slides using the Glycergel mounting media and allowed to dry at room temperature.

**Image acquisition and analysis**

For the representative image of P2YR12, IBA1 and X34 expression at 4 months (Extended Data Fig. 1a), a large field image of the whole sagittal slice was obtained using a Nikon AX Confocal Microscope System driven by NIS-Elements AR (5.41.01) software, using a 4x/0.2 air objective (Nikon CFI Plan Apochromat λ, Nikon # MRD00045). Tiled images were stitched together with an overlap of 15%. For excitation, 405 nm, 488 nm, and 640 nm laser lines were used. Resonant scanning mode was used with 16x line averaging, and minimal crosstalk was set between the channels. Three stacks were acquired using a 5 µm z-interval and images had a resolution of 2048 x 2046, and 1.07 µm pixel size. Higher magnification representative images (Extended Data Fig. 1b) were acquired using a 20x/0.75 air objective (Plan Apo VC 20x DIC N2, Nikon #MRD00201). Twenty-two stacks were acquired using a 0.757 µm z-interval and images had a resolution of 2048 x 2046, and 0.11 µm pixel size.

For the representative image of CD9, HLA-DR, CD68 and X34 expression at 6 months (Extended Data Fig. 1c), images were acquired using an inverted Zeiss LSM 880 microscope with Airyscan detector. The system was equipped with a 20x 1.4 NA Plan-Apochromat objective lens and operated using Zen Black (version 2.3, Carl Zeiss Microscopy GmbH) (0.06 µm pixel size). Ten stacks were acquired using a 1 µm z-interval. Airyscan-acquired images were processed using the default values.

For imaging and analysis of X34, 82E1, LAMP1 and IBA1 after treatment, large field images of the whole slice were obtained using a Nikon AX Confocal Microscope System driven by NIS-Elements AR (5.41.01) software, using a 20x/0.75 air objective (Plan Apo VC 20x DIC N2, Nikon #MRD00201). Tiled images were stitched together with an overlap of 15%. For excitation, 405 nm, 488 nm, 561 nm, and 640 nm laser lines were used and images were acquired using the same acquisition parameters. Three stacks were acquired using a 2.5 µm z-interval. Resonant scanning mode was used with 16x line averaging, and minimal crosstalk was set between the channels. All the quantification results were generated using the measurement of the whole sagittal section. For each experiment, at least 2 sections per mouse were analyzed. For quantification X34+ and 82E1+ areas (1024 x 1024, 0.86 µm pixel size), we used a semi-automated script in Qupath software (v0.4.3)[14]. Briefly, after defining the area of interest (total brain section) with the wand tool, X34+ and 82E1+ areas were identified with the threshold function, and then divided by the total brain section's area. Intensity thresholds were selected manually and used for all the sections, setting the minimum detectable size at 1 µm$^2$ for X34 and 5 µm$^2$ for 82E1. The distribution of X34$^+$ plaques based on area was assessed on high-magnification z-stacks (six cortical images per mouse). Thirty-one stacks were acquired using a Nikon AX Confocal Microscope System controlled by NIS-Elements AR (5.41.01) software, with a 40×/1.25 oil objective (Plan Apo Lambda S 40XC Sil, Nikon #MRD73400) at a resolution of 2048 × 2048 pixels and a pixel size of 0.21 µm. The area of X34+ plaques was analyzed using an automated GA3 recipe in NIS-Elements AR (5.42.05) software. A total of 975, 731, and 942 plaques were analyzed for IgG1, Lecanemab, and Lecanemab LALA-PG, respectively. Plaque distribution was analyzed using a frequency distribution approach, with plaque sizes grouped into intervals of 20 µm to facilitate a structured assessment. The frequency was calculated based on the number of plaques within each bin. The first bin was set to start at 0 µm, and the last bin was set at 2000 µm, ensuring full coverage of observed plaque sizes. While all bins were included in the statistical analysis, only the first 200 µm bins are shown in Fig. 2e, as only a few plaques exceeded this size. LAMP1+ and IBA1+ areas were quantified using an automated GA3 recipe in NIS-Elements AR (5.42.05) software. Images had a resolution of 512 x 512, and 1.71 µm pixel size. Briefly, maximum intensity projections of the Z-stack images were used to identify the section area, using the threshold node. Then, the area of X34+ plaques and IBA1+ area was identified with the threshold node and normalized to the section area; for X34+, minimum detectable size was set at 3 µm. The X34+ area was then expanded of 10 µm, using the circular dilate node. LAMP1+ area was measured in this peri-plaque area; LAMP1 area was then normalized by either the section area or by the peri-plaque area.

For the analysis of synapses, we focused on the cortex and took super-resolution image of X34, Synaptophysin and Homer 1 were acquired using an inverted Zeiss LSM 880 microscope with Airyscan detector in super-resolution mode. At least 4 images (containing at least 2 plaques) were taken per mouse. The system was equipped with a 63x 1.4 NA Plan-Apochromat objective lens and operated using Zen Black (version 2.3, Carl Zeiss Microscopy GmbH). Images (3x3 tiles) were taken with 1.3X electronic magnification (40 nm pixel size). The excitation lasers Argon 488, 514, He-Ne 543, 594 and 633 were used. Airyscan-acquired images were processed using the default values. Obtained images were further processed using Fiji (Imagej-win64). Briefly, a custom script was used to threshold image stacks, expand X34 area of 5 µm and determine density and co-localization of markers. Size exclusion was set between 0.05-1.2 µm.

For imaging OPN+ microglial cells (IBA1+) around X34$^+$ and 82E1$^+$ plaques, confocal fluorescent images were acquired using a Nikon AX Confocal Microscope System driven by NIS-Elements AR (5.41.01) software, using a 40x/1.25 oil objective (Plan Apo Lambda S 40XC Sil, Nikon #MRD73400) (2048 x 2048 resolution, 0.21 µm pixel size). Lasers were set at 405, 488, 561 and 640 nm. Resonant scanning mode was used with 16x line averaging, and minimal crosstalk was set between the channels. Thirty-one stacks were acquired using a 0.5 µm z-interval. All the quantification results were generated on six cortical images per mouse using an automated GA3 recipe in NIS-Elements AR (5.42.05) software. Briefly, maximum intensity projections of the Z-stack images were used to identify the area of X34$^+$ and 82E1$^+$ plaques with the threshold node, with a minimum detectable size set at 3 µm for X34$^+$ plaques. These areas were then expanded of 15 and 5 µm respectively, using the circular dilate node. IBA1$^+$ and IBA1$^+$OPN$^+$ areas were measured in these peri-plaque areas using the threshold node. The ratio of IBA1$^+$/OPN$^+$ area over IBA1+ area was then normalized to the peri-plaque (X34$^+$ and 82E1$^+$) areas.

**Isolation of soluble and insoluble brain extracts and Aβ MSD**

Soluble and insoluble brain extracts were isolated from snap-frozen brain samples. Brain weights were recorded immediately after collection. Ten volumes (w/v) P-TER buffer (Thermo Fisher, #78510) supplemented with cOmplete™ Protease Inhibitor Cocktail (Roche, # 11697498001) and PhosSTOP Phosphatase Inhibitor Cocktail (Roche, #4906845001) were added, and the tissue was homogenized in Lysing Matrix D tube (MP Biomedicals, #6913500) for 45s at 6.5 m/s. Samples were centrifuged for 5 min at 5000 g to remove debris. Subsequently, the supernatant was centrifuged for 1h at 55,000 rpm at 4°C in an Optima Ultracentrifuge using a TLA110 rotor to pellet the insoluble brain fraction. The supernatant (=soluble fraction) was collected and stored at -80°C and the pellet was used for guanidine extraction. Pellets were resuspended in 2µl/mg tissue 6M GuHCl solution (6M GuHCl/50mM Tris-HCl, protease inhibitor cocktail, pH 7.6). Samples were sonicated with a micro-tip for 30 s at 10% amplitude, vortexed for 5 min, and incubated on a shaker for 1h at 25°C and 450 rpm. Samples were ultracentrifuged for 20 min at 70,000 rpm and 4°C. The supernatant, containing guanidine-soluble Aβ fractions (insoluble Aβ) were transferred into a new tube, diluted 12 times with GuHCl diluent (20mM phosphate, 0.4M NaCl, 2mM EDTA, 10% Block Ace, 0.2% BSA, 0.05% NaN3, 0.075% CHAPS, protease inhibitor cocktail, pH 7.0), and stored at -80°C until use.

Aβ38, Aβ40, and Aβ42 levels in the soluble and insoluble brain extracts were quantified by Meso Scale Discovery (MSD). Standard 96-well SECTOR plates (MSD, #L15XA-3) were coated with 0.5 µg ml$^{-1}$ LTDA_38, LDA_40, or LTDA_Aβ42 capture antibodies (homemade mouse monoclonal against Aβ38, Aβ40 or Aβ42 neoepitope respectively) in PBS, pH 7.4 at 4°C overnight. Plates were washed 5 times with PBS with 0.05% Tween 20) and blocked with 0.1 % casein in PBS for 1.5 h at room temperature. Aβ standard curves were prepared with human

Aβ1-38 (rPeptide, #A-1078-1), Aβ1-40 (rPeptide, #A-1153-1) or Aβ1-42 (rPeptide, #A-1163-2). Samples were diluted according to previously determined concentrations. Diluted samples and standards were mixed 1:1 with LTDA_hAβN labelled with a sulfo-TAG detection antibody (250 ng $ml^{-1}$, homemade mouse monoclonal against the N-terminal sequence of human Aβ, in collaboration with Maarten Dewilde), in 0.1% casein in PBS, loaded on the blocked MSD plate, and incubated overnight at 4°C. Plates were washed 5x with PBS-T and 150µl MSD GOLD Read Buffer A (MSD, #R92TG-2) was added to the wells. Plates were read with an MSD Sector Imager 2400A and MESO QuickPlex SQ 120MM (for the experiment performed on immunocompetent mice).

**_In vitro_ plaque clearance assay, image acquisition and analysis**

For the *ex vivo* plaque clearance assay, microglial precursors were collected on day 25 or day 32, and differentiated into microglia-like cells using microglia differentiation medium (TIC; DMEM/F12, glutamine (2 mM), N-acetylcysteine (5 µg $ml^{-1}$), insulin (500 ng $ml^{-1}$), Apo-Transferrin (100 µg $ml^{-1}$), sodium selenite (100 ng $ml^{-1}$), cholesterol (1.5 µg $ml^{-1}$) and heparan sulfate (1 µg $ml^{-1}$) supplemented with 50 ng $ml^{-1}$ interleukin-34, 50 ng $ml^{-1}$ macrophage CSF, 10 ng $ml^{-1}$ CX3CL1 and 25 ng $ml^{-1}$ transforming growth factor-β, based on Abud et al[15]. The medium was changed every other day.

*In vitro* plaque clearance assay was performed as previously described[16–18]. Briefly, 10-µm-thick cryosections from 6-month-old $App^{NL\text{-}G\text{-}F}$ mouse brains[6] were collected onto poly-L-lysine-coated glass coverslips, dried at room temperature for 1 h, followed by incubation with 10 µg $ml^{-1}$ Lecanemab, Lecanemab LALA-PG or IgG1 antibodies in PBS for 1 h at 37°C. Microglia were seeded at $5 \times 10^5$ cells per well in 12-well plates and incubated at 37°C with 5% CO2 for 72 h in microglia differentiation medium. For the osteopontin (OPN) stimulations, we seeded microglia onto the cryosections at $5 \times 10^5$ cells per well and stimulated them with human OPN (Thermo Fisher Scientific, #120-35) at varying concentrations (0, 17, 50, 150, 450, and 1350 ng/mL). The microglia were then incubated at 37°C with 5% $CO_2$ for 72 hours in microglia differentiation medium. For each experiment, sections were either exposed to human microglia or left unexposed as a negative control. After incubation, coverslips were fixed with formaldehyde solution 4% for 15 minutes and washed 3 times with PBS. Afterwards, sections were permeabilized and blocked with PBS containing 0.1% Triton X-100 and 5% donkey serum at room temperature for 1h. Sections were then stained with a mouse anti-amyloid beta (N) (clone 82E1) antibody (0.2 µg $ml^{-1}$; IBL, #10323) overnight at 4°C to visualize Aβ plaques. The next day, sections were washed 3 times with PBS containing 0.1% Triton X (PBS-T) and incubated with donkey anti-mouse secondary antibody, Alexa Fluor™ 647 (2 µg $ml^{-1}$; Invitrogen, #A31571) for 1 h at room temperature, and washed 3 times with PBS-T. Next, sections were blocked with PBS-T containing 5% donkey serum at room temperature for 1h and stained with a biotinylated mouse anti-human CD9 antibody (5 µg $ml^{-1}$; BioLegend, #312112) overnight at 4°C to visualize human microglia. On the next day, sections were washed 3 times with PBS-T and incubated with Streptavidin, Alexa Fluor™ 594 (2 µg $ml^{-1}$; Thermo Scientific, #S32356) for 1 h at room temperature, and washed 3 times with PBS-T. After washing with PBS, coverslips were placed in Mowiol mounting medium on glass slides and allowed to dry at room temperature. Large field images of the whole section were obtained using a Nikon AX Confocal Microscope System driven by NIS-Elements AR (5.41.01) software, using 10x/0.45 air objective (Plan Apo LambdaD - WD 4.0 – Nikon, #MRD70170) with 1.6X electronic magnification (512 x 512 resolution, 2.16 µm pixel size). Tiled images were stitched together with an overlap of 15%. For excitation, 561 nm and 640 nm laser lines were used and images were acquired using the same acquisition parameters. Seven stacks were acquired using a 2 µm z-interval. Resonant scanning mode was used with 16x line averaging, and minimal crosstalk was set between the channels. Quantification of 82E1+ plaque area was performed using an automated General Analysis 3 (GA3) recipe in NIS-Elements AR (5.42.05) software. Briefly, maximum intensity projections of the Z-stack images were used to identify the section area, using the threshold node. Then, the area of 82E1+ plaques was identified with the threshold node and normalized to the section area. All the quantification results were generated using the measurement of the whole sagittal section. For each experiment, 2-4 sections per condition were analyzed.

To confirm that $CD9^+$ cells are also $GFP^+$ and $IBA1^+$, we have also run an experiment using microglia derived from H9 stem cell line stably expressing GFP under the control of chicken β-actin promoter (CAG promoter)[19]. To visualize GFP colocalization with IBA1 and CD9, sections were stained with a mouse anti-amyloid beta (N) (clone 82E1) antibody (0.2 µg $ml^{-1}$; IBL, #10323) overnight at 4°C. The next day, sections were washed and incubated with donkey anti-mouse secondary antibody, Alexa Fluor™ 647 (4 µg $ml^{-1}$; Invitrogen, #A31571) for 1 h at room temperature, and washed 3 times with PBS-T. Next, sections were blocked with PBS-T containing 5% donkey serum at room temperature for 1h and stained with chicken anti-GFP antibody (2 µg $ml^{-1}$; Abcam, #ab13970) combined with biotinylated mouse anti-human CD9 antibody (5 µg $ml^{-1}$; BioLegend, #312112) or rabbit anti-IBA1 antibody (1 µg $ml^{-1}$; Wako, #019-19741) overnight at 4°C. On the next day, sections were washed 3 times with PBS-T and incubated with secondary donkey anti-chicken antibody (2 µg $ml^{-1}$; Jackson, #703-545-155) and Streptavidin, Alexa Fluor™ 594 (2 µg $ml^{-1}$; Thermo Scientific, #S32356) or donkey anti-rabbit, Alexa Fluor™ 594 (2 µg $ml^{-1}$; Invitrogen, #A21207) for 1 h at room temperature, and washed 3 times with PBS-T. After washing with PBS, coverslips were placed in Mowiol mounting medium on glass slides and allowed to dry at room temperature.

**Isolation of human microglia**

Human microglia isolation from the mouse brain was performed as previously described[5]. After perfusion with ice-cold heparinized PBS, one hemisphere (without cerebellum and olfactory bulb) was placed in FACS buffer (PBS containing 2% FCS and 2 mM EDTA) supplemented with 5 µM actinomycin D (ActD; Sigma, # A1410-5MG) for transcriptomics. Brains were mechanically and enzymatically dissociated using Miltenyi Neural Tissue Dissociation Kit P (Miltenyi, #130-092-628) supplemented with 5 µM ActD. Next, samples were passed through a 70-µm strainer (BD2 Falcon), washed in 10 ml of ice-cold FACS buffer with 5 µM ActD and spun at 300 g for 15 min at 4 °C. ActD was kept during collection and enzymatic dissociation of the tissue to prevent artificial activation of human microglia during the procedure[20]. ActD was removed from the myelin removal step to prevent toxicity derived from long-term exposure. Following dissociation, myelin was removed by resuspending pelleted cells in 30% isotonic Percoll (GE Healthcare, #17-5445-02) and centrifuging at 300 g for 15 min at 4 °C. Accumulating layers of myelin and cellular debris were discarded and Fc receptors were blocked in FcR blocking reagents (mouse, 0.1 mg ml$^{-1}$, Miltenyi, #130-092-575; human, 0.1 mg ml$^{-1}$, Miltenyi, # 130-059-901) in cold FACS buffer for 10 min at 4 °C. Next, cells were washed in 5 ml of FACS buffer and pelleted cells were incubated with the following antibodies: PE-Pan-CD11b (20 µg ml$^{-1}$, Miltenyi, #130-113-806), BV421-mCD45 (2 µg ml$^{-1}$, BD Biosciences, #563890), APC-hCD45 (20 µg ml$^{-1}$, BD Biosciences, #555485), Total-Seq A cell hashing antibodies (2 µg ml$^{-1}$, BioLegend) and viability dye (0.5 µg ml$^{-1}$, eFluor 780, Thermo Fisher Scientific, #65-0865-14), in cold FACS buffer during 30 min at 4 °C. After incubation, cells were washed, and the pellet was resuspended in 400 µl of FACS buffer and passed through a 35-µm strainer before sorting. For sorting, the cell suspension was loaded into the input chamber of a MACSQuant Tyto Cartridge, and human cells were sorted based on CD11b and hCD45 expression at 4 °C (MACSQuantify™ Tyto®).

#### ***In vivo* phagocytosis assay**

Starting from 6 months of age, App$^{NL-G-F}$ mice xenotransplanted with human microglia were treated with Lecanemab or IgG1 for 8 weeks. On the day of the experiment, they were injected i.p. with 10mg/kg Methoxy-X04 (Tocris Biosciences, #4920) (protocol adapted from (Lau et al. 2021)) reconstituted in Kolliphor-EL (Sigma, #C5125). Three hours later, mice were euthanized, and microglia was isolated as described above. Cells were washed in 5 ml of FACS buffer and pelleted cells were incubated with the following antibodies: PE-Pan-CD11b (20 µg ml$^{-1}$, Miltenyi, #130-113-806), AF488-mCD45 (2 µg ml$^{-1}$, Biolegend #109815), APC-hCD45 (2 µg ml$^{-1}$, BD Biosciences, #555485), and viability dye (0.5 µg ml$^{-1}$, eFluor 780, Thermo Fisher Scientific, #65-0865-14), in cold FACS buffer for 30 min at 4 °C. After incubation, cells were washed and fixed with eBioscience Foxp3 Fixation/Permeabilization kit (eBioscience, #00-5521-00) for 30 min at 4 °C. For intracellular CD68 staining, fixed cells were permeabilized (eBioscience, #00-8333-56), washed and incubated in PE Vio615 CD68 (10 µg ml$^{-1}$, Miltenyi, #130-114-656) in permeabilization buffer overnight at 4 °C. Cells were washed with FACS buffer, the pellet was resuspended in 500 µl of FACS buffer and passed through a 35-µm strainer prior to FACS acquisition. Flow data was acquired on a BD Fortessa (BD FACSDiva Software, version 9.7). Dead cells and doublets were gated out prior to downstream analysis. Human microglia were identified by gating on hCD45. Methoxy-x04 gating was set with negative controls (xenografted App$^{Hu}$ mice injected with Methoxy-x04). Methoxy-x04 populations within the hCD45 and within the CD68-positive gate were analysed for Methoxy-X04 incorporation. FACS data was analysed with FlowJo software (v10.8.1).

#### **Single-cell libraries preparation and sequencing**

For single-cell RNA sequencing, 15,000–20,000 human microglia (CD11b+, hCD45+) from each mouse were sorted on the MACSQuant Tyto and diluted to a final concentration of 1,000 cells µl$^{-1}$. Since all the samples were individually hashed using Total-Seq A cell hashing antibodies, 2,000-4,000 human microglia per animal were pooled and loaded onto the Chromium Next GEM Chip G (PN no. 2000177). The DNA library preparations were generated following the manufacturer's instructions (CG000204 Chromium Next GEM Single Cell 3′ Reagent Kits v3.1). In parallel, the hashtag oligo libraries were prepared according to the manufacturer's instructions (BioLegend, Total-Seq A Antibodies and Cell Hashing with 10x Single Cell 3′

Reagent Kit v3 3.1 Protocol) using 16 cycles for the index PCR. A total of 5 libraries containing 12 biological replicates were sequenced (GAL002, GAL004, GAL005, H1 and H2, Extended Data Figs. 4a-b), targeting a 90% messenger RNA and 10% hashtag oligo library (50,000 reads per cell), on a HiSeq4000 (Illumina) platform with the recommended read lengths by the 10X Genomics workflow.

### Single-cell sequencing data analysis

#### Alignment, pre-processing, and quality control

The 10x Genomics Cell Ranger software (v6.1.2) was used to align reads to a combined human/mouse reference genome (GRCh38 and mm10), demultiplex cellular barcodes, and quantify unique molecular identifiers (UMI) and Hashtag oligos (HTO). The UMI and hashtag count matrices for each of the libraries were loaded into the Seurat package (v4.1.1, R)[21]. Genes whose transcripts were identified in less than 3 cells, as well as cells with fewer than 100 unique transcripts, were removed from the expression matrices. Demultiplexing of cells to their sample of origin was performed as described in Stoeckius *et al.*[22]. In brief, cell barcodes were filtered to include only those detected in both RNA and HTO data. The HTO assays were normalized using the centered log-ratio (CLR) transformation. Seurat's *HTODemux()* function was run with default parameters on the hashtag-count matrices to demultiplex the cells. Cells that could not be assigned to an HTO and cells that were positive for more than one HTO were deemed as negatives and inter-sample doublets, respectively, and removed from further analysis. In the remaining demultiplexed singlets (n=27,290), putative low-quality cells were identified by setting a threshold of 2.5 median absolute deviations (MADs) on the QC covariates of interest per sequenced library. These covariates are: log1p(number of transcripts detected), log1p(number of genes detected), and percentage of transcripts mapping to human mitochondrial genes. Furthermore, any cell with more than 5% of reads aligned to the mouse transcriptome were removed. Finally, intra-sample doublets were identified in each library using DoubletFinder (v2.0.3) and subsequently removed. The remaining high-quality single-cell transcriptomes (n=22,841) were used for further analysis.

#### Normalization, integration, and clustering

Gene counts from each of the 5 libraries were individually normalized using *SCTransform()*[23]. The *SelectIntegrationFeatures()* function was used to identify the 3,000 most variable features in each library for downstream analysis. To integrate the libraries, we ran *PrepSCTIntegration()*, followed by the *FindIntegrationAnchors()* and the *IntegrateData()* functions. A principal component analysis (PCA) was performed on the integrated expression matrix with *RunPCA()*, after which Uniform Manifold Approximation and Projection (UMAP)[24] embeddings were created with *RunUMAP()*, using the first 30 principal components (PCs) as input. A shared nearest neighbor (SNN) graph was constructed using the *FindNeighbors()* function, again using 30 PCs. Unbiased clustering by the Louvain algorithm (*FindClusters()*) with a resolution of 0.35 identified 11 clusters (Extended Fig. 4b,c). The *PrepSCTFindMarkers()* and *FindAllMarkers()* functions were run to find the marker genes for each cluster. Cluster 10 was identified as the macrophage cluster and removed from analysis (Extended Data Figs. 4c,d). The above-described steps were repeated for the remaining microglial cells (n= 22,420) using the same parameters, except for the unbiased Louvain clustering, which was run with a resolution of 0.15 and identified

9 clusters. Cell state identities were assigned to each of the clusters based on the expression of marker genes, as determined in "Differential Gene Expression Analysis" and visualized by Seurat's *DotPlot()* (Extended Data Fig. 4f).

**Single-cell Differential Gene Expression Analysis**

Differential expressions between clusters and treatment conditions were performed on the SCT-normalized counts using the *PrepSCTFindMarkers()* and *FindMarkers()* functions provided within the Seurat package. P-values were calculated using the Wilcoxon rank-sum test and were corrected for multiple testing using the Bonferroni method.

**Differential abundance assessment of microglia cell states**

To test for differential abundance of cell states in association with Lecanemab treatment, we used Mixed-effects modeling Association of Single Cells (MASC)[25] (v0.1.0, R), which assesses the association between a cell's cluster identity and its condition status (Extended Data Fig. 4c). We included treatment (Lecanemab or IgG1) as a fixed effect, while the mouse ID was included as a random effect.

**Gene Set Enrichment Analyses**

Gene Set Enrichment Analyses (GSEA)[26] was performed using the clusterProfiler[27] package (v4.6.2). The outputs of differential expression analyses between conditions (in the scRNA-seq analyses) or in function of distance to pathology (in the Nova-ST data) were ranked according to each gene's log2(FoldChange). ClusterProfiler's *gseKEGG()* and *GSEA()* functions were run on this ranked list to calculate gene set enrichment scores for KEGG pathways (, Figs. 1f-g, Extended Data Figs. 2d-e) and WGCNA modules (Fig. 4d, Fig. 5b, Extended Data Fig. 5e), while *gsePathway()* from the ReactomePA[28] (v1.42.0, R) package was run to do the same for Reactome pathways (Extended Data Fig. 4e). The following parameters were used: *nPerm* = 1000, *minGSSize* = 40, *maxGSSize* = 800, *pvalueCutoff* = 0.05, qvalueCutoff = 0.05, *pAdjustMethod* = "BH". The GSEA enrichment profiles of selected signatures were plotted using enrichr's gseaplot2() function.

**Weighted Gene Co-expression Network Analysis (WGCNA)**

To perform co-expression network analysis with WGCNA[29], we employed High-Definition Weighted Gene Co-expression Network Analysis (hdWGCNA)[30] (v0.3.0, R), specifically designed for analysis of high dimensional snRNA-seq data. Briefly, this approach groups transcriptionally similar cells into "metacells", so as to overcome the issue of sparsity of single cell data and thus providing more robust gene-gene correlation estimates. First, we filtered out genes expressed in less than 5% of all cells. We next generated metacells per sample with the *MetacellsByGroups()* function which employs a k-Nearest Neighbors (KNN) algorithm, using the parameters *target_metacells*=200, *group.by*="sample", *reduction*="pca", *k*=25, *max.shared*=10. Next, we used the *TestSoftPowers()* function to determine the optimal soft power threshold for constructing a "signed hybrid" co-expression network. We set the minimum soft power threshold at the minimum value with a Scale Free Topology Model Fit of at least 0.9, so as to retain robust gene-gene correlations while eliminating weak links in the adjacency matrix. Finally, we performed a signed-hybrid network construction and module detection with the *ConstructNetwork()* function. The module dendrogam was visualized using the *PlotDendrogram*() function (Extended Data Fig. 5a). The *ModuleEigengenes()* function was run to determine the module eigengenes (MEs), or the first principal components of the expression matrix specific to each of the modules. The intra-modular connectivity (kME) of each gene, representing the correlation of the gene with its ME, was calculated using *SignedKME*(). The 10 genes with the highest kMEs per module were identified as hub genes and their connectivities were plotted using the ggraph and tidygraph packages (v2.1.0 and v1.2.3, respectively, R) (Fig. 4e).

**Gene Set Enrichment Visualization**

To calculate and visualize each cell's enrichment of gene sets of interest (such as markers for microglial states and WGCNA modules), we used Seurat's *AddModuleScore()* function. In brief, this function calculates the activity of a gene set in each cell by comparing the mean abundance level of the genes of interest against the average abundance of sets of random control genes which have a similar average expression level. The calculated scores were visualized on the UMAP (Extended Data Fig. 5c).

**Gene Set Overrepresentation Analyses**

Over-representation of Gene Ontology[31,32] terms (Biological Processes and Molecular Functions) in the WGCNA modules was assessed with clusterProfiler's *enrichGO()* function (Fig. 4e, Table 2). The analyses were run using the parameters: *minGSSize* = 10, *maxGSSize* = 500, *pvalueCutoff* = 0.05, *qvalueCutoff* = 0.2, *pAdjustMethod* = "BH". Overlap of each of the modules with the ATM[33] and PAM[34] gene sets was assessed using a hypergeometric test, implemented through the stat package's (v.4.5.0, R) *phyper*() function (Fig. 5a, Table 3). Resultant p-values were corrected for multiple testing using the Benjamini–Hochberg method.

**Statistical analysis**

Statistical analyses and data visualization were performed using GraphPad Prism (v9) and R (v4.2.3). Data are presented as scatter dot plot with bar, and the line at the mean ± SEM (standard error of the mean). All n values represent individual animals, unless stated otherwise (i.e., *ex vivo* plaque clearance assay, where each n represent an independent experiment). When appropriate, animals were randomly assigned to conditions and conditions were randomized to account for potential ordering effects. To avoid litter bias in the mouse experiments, experimental groups were composed of animals from different litters randomly distributed. All analyses were conducted either blindly to the experimental condition or using an automated GA3 recipe in NIS-Elements AR software. For all the ELISA data, statistical outliers (caused by technical errors) were identified using the ROUT test in Prism10 (Q = 1%) and excluded from further analysis. Normality of residuals was checked with the Shapiro–Wilk tests. Comparisons between two groups following a normal distribution were analyzed using two-tailed unpaired t-test, comparisons between two groups not following a normal distribution with Mann-Whitney test. When three groups were compared and data were normally distributed, ordinary one-way ANOVA was used; when significant, it was followed by Bonferroni's multiple comparisons test. When data were not normally distributed, ranks were compared with Kruskal-Wallis followed by Dunn's multiple comparisons test. To determine the statistical significance of the difference between microglia and no-microglia curves upon OPN stimulation, we used a modified Chi-squared method (as described in [35]). Notably, this method includes a correction for the deviation from normality. To determine the overall statistical significance of the differences in plaque area distribution, we used the Anderson-Darling test. For pairwise comparisons between distributions, we performed the Kolmogorov-Smirnov test. To account for multiple comparisons (three in total), we applied a Bonferroni correction by multiplying the obtained p-values by three. The statistical tests are reported in the figure legends and for all analyses, $\alpha = 0.05$. A detailed description of statistical analysis and number of mice used in this study is reported in **Table S2**. For single-cell sequencing, data generated in this study are available at Gene Expression Omnibus (GEO) database with accession number that will be provided upon publication. Single cell and Nova-ST analyses will be made available at: https://github.com/mmzielonka/AlbertiniZielonka_Lecanemab_2025.git. Other data are available upon request.

**Table S1 Summary of antibody sequences used in this study.**

**Table S2 Detailed description of statistical analysis used in this study.**

**Supplementary Movie 1 3D Surface Reconstruction of Lecanemab Localization in Microglia.** This video presents a 3D surface reconstruction of D54D2-positive plaques in a Lecanemab-treated mouse, generated using Imaris surface reconstruction. The anti-IgG signal (labeling Lecanemab) is detected within human $CD45^{+}$ microglia, highlighting its localization inside microglial surfaces. This reconstruction corresponds to Fig. 1a, upper panel**.**
